# Cellular senescence is associated with age-related loss of liver zonation and hepatocyte function

**DOI:** 10.64898/2026.08.08.743614

**Authors:** Linshan Laux, Alisha Aristel, Syed Ali, Kathryn Lande, Meiyi Li, K. Garrett Evensen, Aaron Havas, Zhen Miao, Jane Zhang, Samuel Peters, Jiayi Hu, Luise Angelini, Maggie Klaers, Jennie Brocksome, Adam Lewis, Nishitha Paidimukkala, Mary E. Brown, Chase M. Carver, Marissa J. Schafer, Jeffrey H. Albrecht, April Wehner, Peter Adams, Constantin Aliferis, Oyedele Adeyi, Sundeep Khosla, Xiao Dong, Jinhua Wang, Paul D. Robbins, Nancy Zhang, Laura J. Niedernhofer

## Abstract

The liver is organized into tightly regulated zones with distinct metabolic functions but zonation erodes with age. Cellular senescence contributes to aging and liver diseases, however, its impact on aging biology is ill-defined. As part of The Cellular Senescence Network Consortium, we used multiple spatial transcriptomics approaches (GeoMx, Visium, CosMx) with snRNA-seq to profile senescence signatures, zonation markers, and metabolic pathways in livers from wild-type (WT) mice of multiple ages. We observed a loss of canonical zone signatures in aged mouse livers characterized by “expansion” of midlobular (zone 2) marker gene expression, accompanied by diminished expression of zone 3 marker genes by middle-age (18 months), indicative of loss of cell identity. Multiple analytic approaches identified distinct age-, zone- and sex-specific senescence signatures, which were significantly associated with zonation markers changes. This was recapitulated in *Ercc1* mutant models of accelerated senescence, supporting a causal role of senescent cells in liver aging. A “no-zone” hepatocyte-like cluster expanded with age and with the strongest Senescence-Associated Secretory Phenotype (SASP) profile. Gene expression profiles from senescent hepatocytes implicate decreased WNT signaling and increased BMP as contributing to age-related loss of zonation. Together, these data elucidate the role of senescent cells in driving aging biology in non-diseased liver through disruption of cell:cell signaling and the loss of metabolic and cell identity gene expression necessary for hepatocyte function.

## Introduction

Aging is associated with morphological, molecular and functional changes of the liver[1–3]. Cellular senescence has been identified in different cell types in the liver[4, 5] and associated with acute and chronic liver injury and disease[6–8]. Senescent hepatocytes undergo morphological changes and metabolic rewiring leading to altered liver function and disease progression[5, 9]. Liver sinusoidal endothelial cells are one of the prominent senescent liver cell types that contributes to both structural and functional changes[4, 10]. Hepatic stellate cells senescence is associated with liver regeneration, fibrosis regression and progression dependent on disease status[11]. Senescent cholangiocytes are also associated with liver fibrosis and cholangiopathy[12, 13]. In addition, distinct roles of senescent macrophage and endothelial cell populations were identified in liver regeneration using *p16^Ink4a^* tracer mice[4]. Most recently, a p21^+^TREM2^+^ senescent macrophages population was identified to be a central driver of inflammaging and metabolic liver diseases[14].

The liver consists of hexagonal shaped lobule units with the external vertices defined by portal triads and the center by a single central vein. There is a gradation of metabolic functions along the radial axis of liver lobules defined as zones 1-3 from external to internal[15]. Numerous studies established that hepatocytes from different zones are more susceptible to different pathologic stimuli, demonstrating their divergent cell identities[16, 17]. Master regulatory pathways of liver zonation have been extensively investigated including the Wnt/β-catenin as well as the R-spondin (RSPO)-LGR4/5 axis[18–22]. Notch, Insulin/glucagon signaling and the oxygen gradient across the lobule also play crucial roles in liver zonation and distinct metabolic activities[23, 24]. In the pericentral zone 3, endothelial cells play a critical role in the dynamic control of liver zonation and regeneration by secreting ligands WNT2 and WNT9b[25]. Midlobular zone 2 hepatocytes are the major contributor to maintaining liver homeostasis and hepatocyte replenishment[26], which is driven by insulin-like growth factor binding protein 2-mechanistic target of rapamycin-cyclin D1 (IGFBP2-mTOR-CCND1) signaling[27].

Aging alters mechanical properties of liver, for example, increased liver stiffness due to extracellular matrix (ECM) deposition. Zone 2 hepatocytes are sensitive to biomechanical forces activating the mechnotransduction DPP4+-PIEZO1-IGFBP2 axis to drive regeneration across liver zones[28]. In addition, the transforming growth factor-beta (TGF-β) superfamily including the bone morphogenetic proteins (BMPs) branch which activates SMAD1/5/8 signaling and non-SMAD signaling are implicated in liver homeostasis and pathophysiology[29, 30]. BMPs regulate hepatic iron metabolism by promoting the expression of hormone hepcidin (encoded by *Hamp*)[31]. BMPs are also well-established senescence-associated secretory phenotype (SASP) factors[32]. How aging and in particular, cellular senescence alters liver zonation and function remains to be further explored.

In this study, we used single nuclei and multiple spatial transcriptomics platforms to investigate age-related changes in liver of wild-type (WT) mice across lifespan and murine models of accelerated systemic and endothelial-specific senescence[33] to interrogate this gap in knowledge. We found that senescence impacts the identity, metabolic function, cell signaling, and regenerative capacity of hepatocytes during aging.

## Results

### Loss of canonical zone signatures in aged mouse livers

It is well established that liver zonation undergoes dynamic changes upon physiological and pathological demands, including aging. However, defining how the zonation pattern changes with age and the mechanism behind it requires further investigation[34, 35]. To gain a thorough characterization of zone signatures, we used multiple spatial transcriptomics approaches to profile zonation marker genes in livers from wild-type (WT) mice across multiple ages. Immunofluorescence staining utilizing three zonation demarcation markers revealed a well-organized hexagonal zonation pattern in both young female and male WT mouse livers (Figure 1a). However, a clear disruption of organization was observed in livers of mice >2-years-of-age, characterized by expression of midlobular (zone 2) marker hepcidin+hepcidin2 ‘invading’ into neighboring zones, in particular pericentral (zone 3) regions. This was accompanied by disorganized expression of the periportal (zone 1) marker E-cadherin. Zone 3 appeared the most disrupted as the central veins were surrounded by hepatocytes often expressing both zone 3 and zone 2 markers in old animals (Figure 1a).

**Figure 1.**
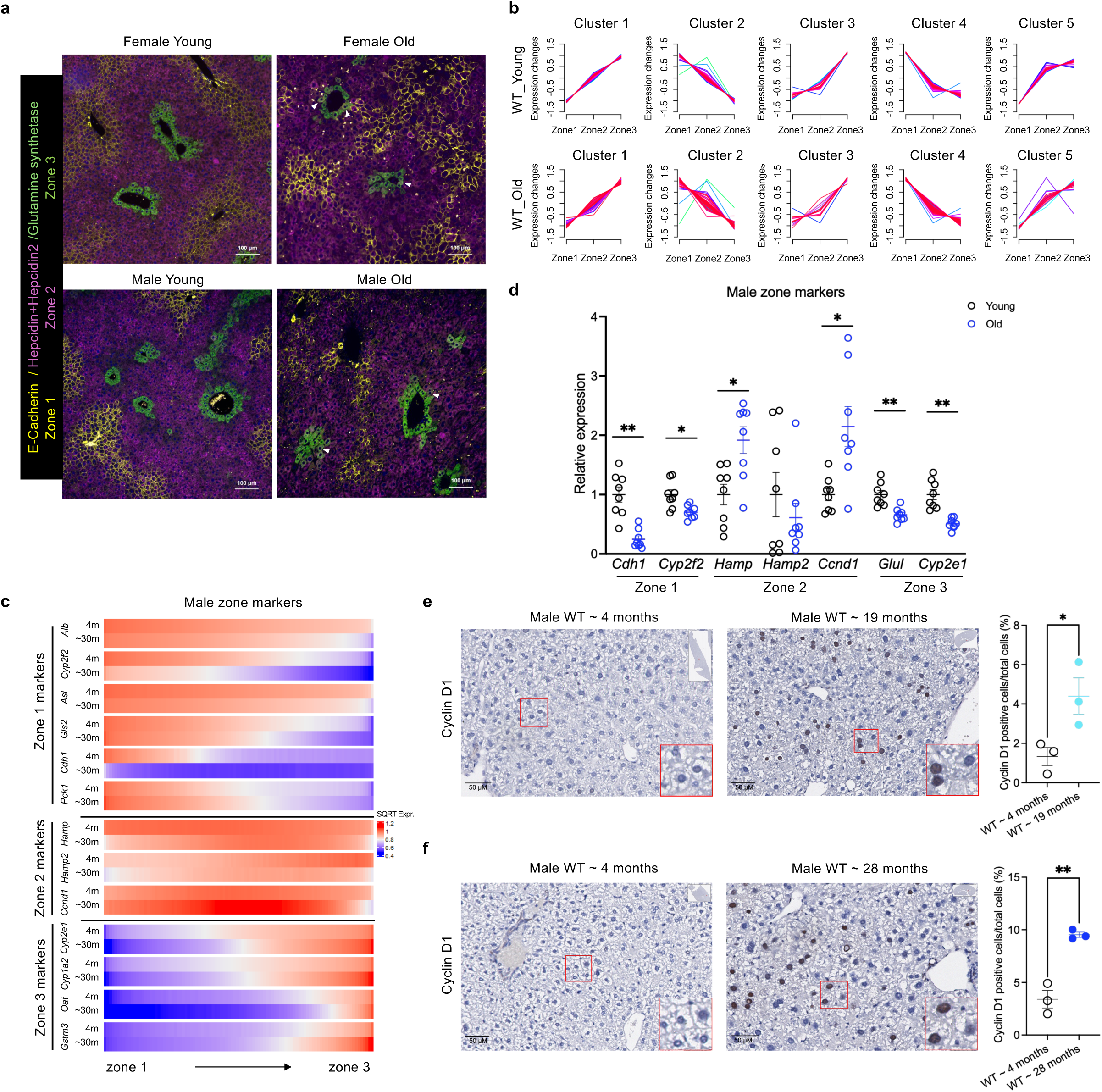
Age-related disruption of liver zonation identified by multiple methods. a. Immunofluorescent staining of zonation markers: E-Cadherin (yellow): periportal (zone 1), Hepcidin+Hepcidin2 (magenta): midlobular (zone 2), Glutamine synthetase (green): pericentral (zone 3). White arrowheads pointing to hepatocytes expressing both zone 2 and zone 3 markers. b. GeoMx digital spatial profiling identified 215 genes differentially expressed across the three hepatic zones in young WT moue livers (top panel). These aligned into 5 clusters based on differing expression patterns across liver zones. Each line represents the expression variance of a single gene across the three zones. Warm colors (red and pink) indicate high confidence with which the gene belongs to that cluster, whereas cool colors (blue and green) indicate lower confidence. The same 215 genes were analyzed in WT old livers, and their expression patterns were illustrated in the bottom panel. Gene expression changes were most disrupted with age in zone 2 across all 5 clusters. c. Heatmaps showing the gene expression pattern of zone markers (y-axis for each gene analyzed) across the 3 liver zones (x-axis) in young (4-month-old)) and old (∼ 30-month-old) male mouse livers measured by Visium transcriptomics studies. d. qPCR analysis of zone marker gene expression in young and old male mouse livers. Each circle represents an individual mouse (n=8 per age group). Multiple unpaired *t*-tests and adjusted *p* values are presented as: \**p* < 0.05, \*\**p* < 0.01. e. Representative images of immunohistochemical staining for the midlobular zone marker Cyclin D1 in young (∼4-month-old) and middle age (∼19-month-old) male mouse livers, stained on the same slide. Two-tailed student *t*-test was applied for Cyclin D1 positive cell (%) quantification comparison, \**p* < 0.05. n=3 for each age group. Red box is magnified in the bottom right corner of the panel to illustrate staining. f. Representative images of immunohistochemical staining for the midlobular zone marker Cyclin D1 in young (∼4-month-old) and old (∼28-month-old) male mouse livers, stained on the same slide. Two-tailed student *t*-test was applied for Cyclin D1 positive cell (%) quantification comparison, \*\**p* < 0.01. n=3 for each age group. Red box is magnified in the bottom right corner of the panel to illustrate staining.

To thoroughly characterize age-associated zonation changes, spatial transcriptomics was used. On Bruker/NanoString GeoMx digital spatial profiler, regions of interest (ROIs) were selected and zonation manually defined in each ROI, within which whole transcriptome profiling targeted at all cell types but not single cells (Extended Data Figure 1a). To identify reliable zonation markers, we identified 215 differentially expressed genes (DEGs) across the three liver zones (Supplementary Table 1) in GeoMx profiling of young WT mouse livers. Principle component analysis (PCA) revealed clustering by zone rather than age, suggesting the DEGs captured distinct features of different zones (Extended Data Figure 1b).

To further explore how shifts in gene expression contribute to zone-specific functions, we analyzed their expression patterns across the three zones and categorized them into 5 clusters in the WT young livers (Supplemental Figure 1c, top panel). Genes in clusters 1 and 2 showed gradual upregulation or downregulation from zone 1 to zone 3, respectively; genes in cluster 3 were upregulated only in zone 3; genes in clusters 4 and 5 were significantly upregulated or downregulated only in zone 1, respectively. KEGG pathway enrichment analysis for each cluster identified significant enrichment of a total of 40 KEGG pathways (Supplemental Figure 1c, lower panel). The genes upregulated in zone 1 in young livers (clusters 2 and 4) were enriched in gluconeogenesis, amino acid metabolism, and AMPK signaling pathways, suggesting active energy metabolism in this zone, consistent with its proximity to the portal vein with high oxygen and nutrient availability. The genes upregulated in zone 3 (cluster 1, 3 and 5) were associated with metabolism of retinol, xenobiotics, fatty acid (particularly arachidonic and linoleic acids) and cholesterol, which is consistent with the role of zone 3 in detoxification and drug metabolism. To study whether aging or senescence influences the expression of the zone-specific DEGs, we assessed the same genes from each cluster in the old WT livers and analyzed their expression patterns. We observed that with age, expression of zone 2 zonation genes changed most dramatically, suggesting that genes in this zone might be especially sensitive to aging or senescence-related changes (Figure 1b).

Visium transcriptomics, which carries out whole transcriptomics profiling at spot level (multiple types of cells) but not at single cell level, was also used to profile expression changes in zonation markers in multiple age groups of female mice, as well as young and old male mouse livers. Consistent with the immunohistochemical data, we observed expansion of midlobular (zone 2) marker gene expression, particularly *Hamp* and *Hamp2* in female (Extended Data Figure 1d) and *Ccnd1* in male (Figure 1c), along with diminished expression of the zone 3 and zone 1 marker genes. Such zone changes are progressive and already obvious by middle-age (18 months) in female livers (Extended Data Figure 1d). We also observed drastic differential gene expression patterns between female and male zone marker genes, indicating sex-related differences (Figure 1c vs. Extended Data Figure 1d).

qPCR confirmed zone marker gene mRNA level changes, showing decreased expression of zone 3 and zone 1 markers (Figure 1d; Extended Data Figure 1e) with increased expression of zone 2 genes in old WT livers (Figure 1d). Immunohistochemistry staining further confirmed increased zone 2 marker Cyclin D1 signal in ∼19-month-old and ∼28-month-old WT male livers relative to 4-month-old animals (Figure 1e, f). Overall, all the above data reveal a disrupted liver zonation pattern associated with aging.

### Age- and zone-related differentially expressed genes identified across the liver lobule by Visium spatial transcriptomics

Upon identifying age-related zonation changes, we investigated age-related DEGs across liver lobule using Visium spatial transcriptomics (Figure 2a). We profiled the spatial expression pattern of the DEGs across liver zones in female and male livers with aging (Figure 2b). Pathway analysis suggested the upregulated age-related DEGs were involved in lipid metabolism, chemotaxis, angiogenesis, cell adhesion, inflammatory response and cellular senescence; accompanied by down-regulation of WNT signaling, cell growth and proliferation, and various hepatic metabolic activities compared to the young samples (Figure 2c). Notably, >40% of the age-related DEGs that were increased in old female and male livers encode secreted proteins or extracellular-associated factors (Figure 2d).

**Figure 2.**
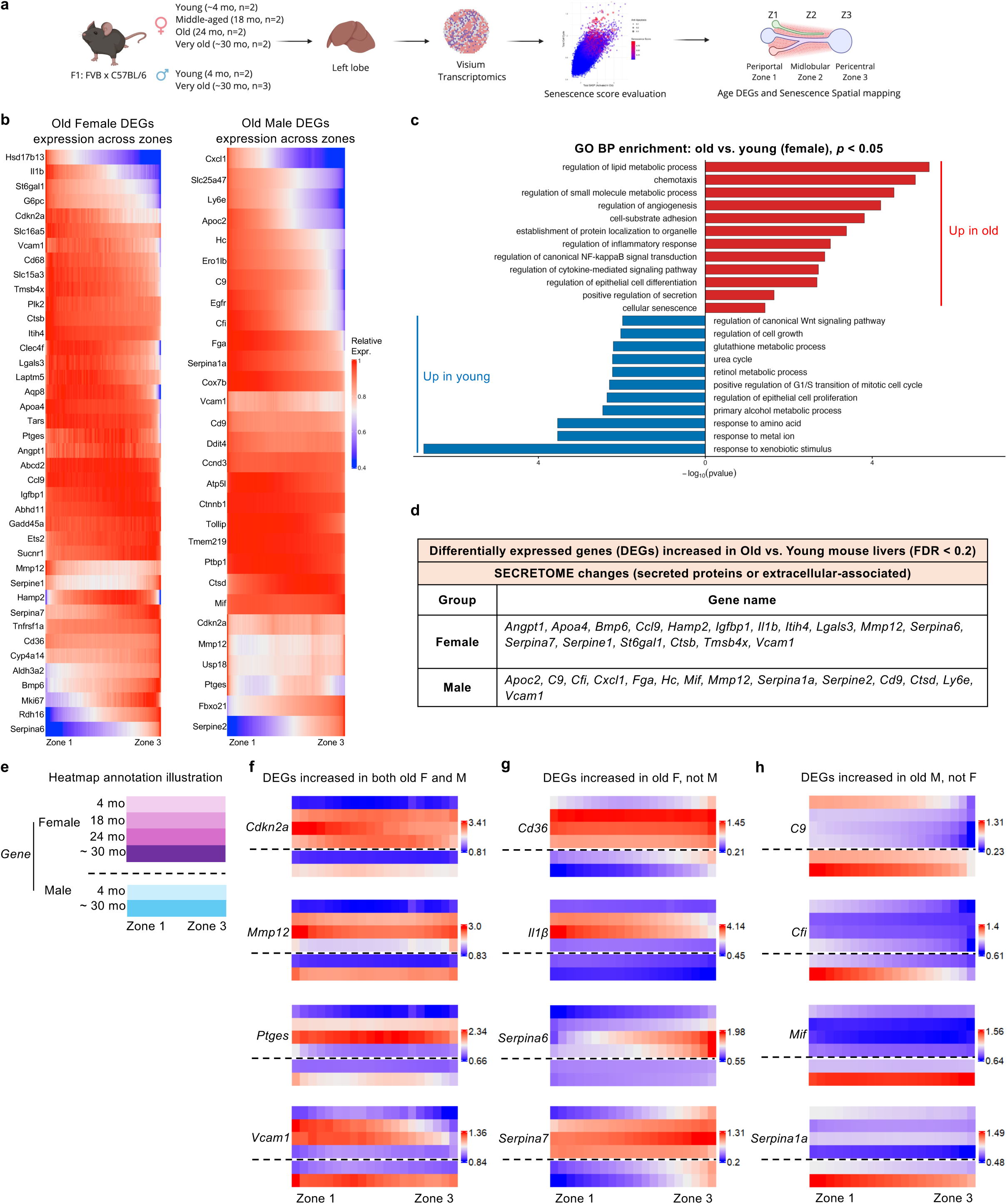
Age- and zone-related differentially expressed genes identified across the liver lobule by Visium spatial transcriptomics. a. Schematic illustration of the experimental design of the Visium transcriptomics study. b. Heatmap of zone-related differentially expressed genes (DEGs, y-axis) identified in 24-month-old female livers (left) and 30-month-old male livers (right) across liver zones (x-axis) compared with the respective young controls (4-month-old WT mice). DEGs were identified using limma analysis, with FDR < 0.2 defined as significant. c. Gene Ontology Biological Process (GO BP) enrichment analysis was performed on the significant DEGs identified by limma analysis between old (24-months-old) and young (4-months-old) female WT mouse livers (FDR < 0.2). Enriched GO terms with *p* < 0.05 are shown. Pathways on the right (red) are enriched among genes upregulated in old livers, whereas pathways on the left (blue) are enriched among genes upregulated in young livers. d. Table of age- and zone-related significant DEGs across liver zones that encode secreted or extracellular-associated proteins identified significantly increased in the old vs. young mouse livers. The DEGs were identified using limma analysis with FDR <0.2 defined as significant. e. Illustration of how the heatmaps in f-h are organized. Each row in the heatmaps represents a different age group of female mice (top 4 rows) and male mice (bottom 2 rows). On the x-axis, DEG expression is indicated across the liver lobule from zone 1 (left) to zone 3 (right). f. Heatmaps illustrating the expression pattern of four common age- and senescence-related DEGs significantly upregulated in livers of both old female and male mice across liver zones. g. Heatmaps illustrating the expression pattern of representative age-related DEGs significantly upregulated in livers of old female mice but not males across liver zones. h. Heatmaps illustrating the expression pattern of representative age-related DEGs significantly upregulated in livers of old male mice but not females across liver zones.

Many well-established senescence genes were identified in these age- and zone-related DEGs. Four senescence-associated genes were significantly upregulated in both old female and male livers (e.g., *Cdkn2a*, *Mmp12, Ptges and Vcam1*) (Figure 2e, f). Some DEGs increased with age only in females, for example, *Cd36[36]* and *Il-1β* (Figure 2g); or only in males, for example, *C9* and *Mif[32]* (Figure 2h). Significantly increased expression of senescence genes such as *Cdkn2a, Cdkn1a, and Timp1* in aged WT mouse livers (>2-years-old) were confirmed by qPCR (Extended Data Figure 2a). Overall, the identified age- and zone-related DEGs were enriched in up-regulated genes related to immune response, cytokine signaling and extracellular matrix remodeling, and genes with reduced expression related to metabolic and mitochondrial function, all closely related to cellular senescence.

### Age-, sex- and zone-related senescence signatures identification by Visium spatial transcriptomics

To further spatially identify senescence across the liver lobule, we applied three categories of senescence-associated genes (SASP, cell cycle inhibitor, anti-apoptosis) (Supplementary Table 2) to create a “senescence score” with additional *Mki67* filtering (see Methods) to identify senescent pixels in Visium spatial transcriptomics profiled mouse livers (Figure 2a). Four prominent features were identified in our analysis. First, compared to the young animals, there was a significantly higher number of pixels with high senescence scores in old female (Figure 3a) and male mouse livers (Figure 3b), consistent across all samples. Second, the increase in senescence was detected as early as 18-months-of-age, increased by 24-months but was decreased in 30-month-old female. The higher senescence score observed in 18- and 24-months livers were driven by increased expression of SASP components (Figure 3a). Notably, this is consistent with the time course of zonation changes in female mouse livers (Extended Data Figure 1d). Third, high senescence scores were identified across the entire liver lobule with a heterogeneous distribution pattern in different zones (Figure 3c and g; Extended Data Figure 3a and e). Lastly, pixels with a high senescence score appear to cluster with age especially at advanced age (Figure 3d-f and h-j; Extended Data Figure 3b-d and f-h).

**Figure 3.**
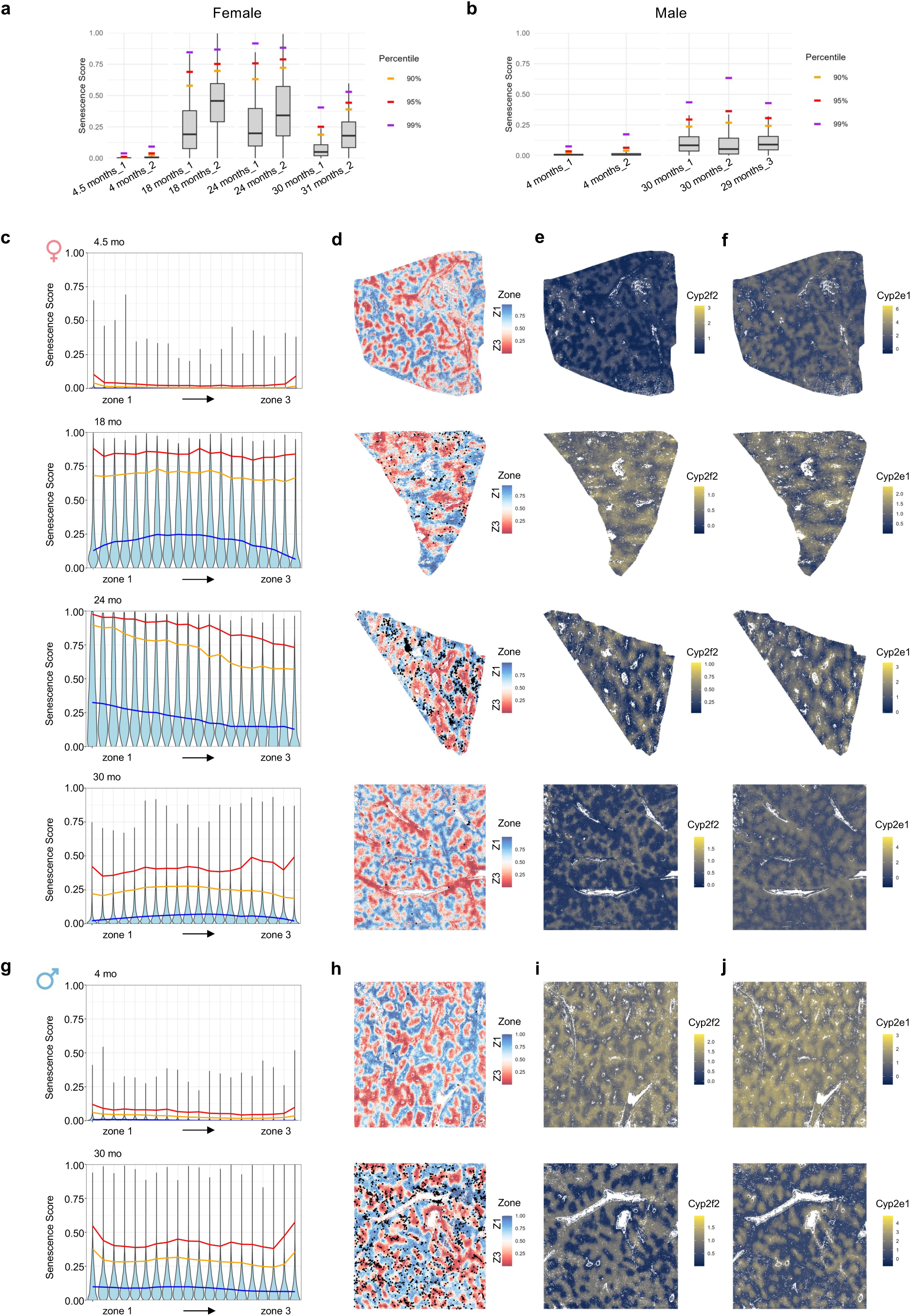
Age-, sex- and zone-related senescence signature identification by Visium spatial transcriptomics. a. Boxplots show the distribution of pixel-level senescence scores within each individual mouse liver sample. The score was calculated for each spatial pixel using SASP, cell cycle inhibitor, and anti-apoptosis gene modules with *Mki67* filtering applied to mask out pixels with *Mki67* expression level in the top 95^th^ percentile. Percentile markers indicate the 90^th^, 95^th^, and 99^th^ percentiles of the pixel-level senescence score distribution for each sample. Statistical testing was performed at the pixel level with Wilcoxon-rank sum test comparing senescence score distributions between age groups, which yielded a significant increase in 18, 24 and 30-month-old samples compared to the young (*p* value < 10^-16). b. Same as a. except for male mice (*p* value < 10^-16). c. Violin plots showing senescence scores across the liver lobule from zone 1 to zone 3 (x-axis) in different age groups (4.5, 18, 24, 30-months old, by panel top to bottom) of WT female mice. Liver zone was divided into 20 bins from zone 1 to zone 3, and each violin line represents the distribution of senescence score for each zonation bin. Blue line represents 50^th^ percentile, orange: 95^th^ percentile, and red: 99^th^ percentile. d. Spatial mapping of senescence-high pixels (black dots represent the top 98^th^ percentile of senescence score pixels in 24-month-old samples at a cutoff threshold of 0.86 [in Figure 3a]) in the zones of WT female mouse livers. Zone 1 blue, zone 3 red, at 4.5, 18, 24 and 30-months of age (top to bottom). e. Spatial distribution of gene expression of the zone 1 marker *Cyp2f2* in WT female mouse livers at 4.5, 18, 24 and 30-months of age (top to bottom). f. Spatial distribution of gene expression of the zone 3 marker *Cyp2e1* in WT female mouse livers at 4.5, 18, 24 and 30-months of age (top to bottom). g. Same as c, but for WT male mice at 4 and 30-months of age. h. Same as d, but for WT male mice at 4 and 30-months of age. (Black dots represent the top 98^th^ percentile of senescence score pixels in 30-month-old samples at a cutoff threshold of 0.41 [in Figure 3b]). i. Same as e, but for WT male mice at 4 and 30-months of age. j. Same as f, but for WT male mice at 4 and 30-months of age.

After identifying high senescence score pixels using the targeted approach, we investigated DEGs that were significantly correlated with these senescent pixels (Figure 4a and c). Three clusters of senescence-associated genes were identified across female (Figure 4a) and male livers (Figure 4c). Cluster 1 positively associated with senescence (Supplementary Table 3). Pathway analysis of Cluster 1 genes revealed enrichment of cytokine, integrin signaling, ECM-receptor interaction, complement and coagulation cascades, and numerous metabolism related pathways in female (Figure 4b) and male livers (Figure 4d), suggesting profound functional effects linked to senescence. This approach allowed us to also identify up-regulated genes significantly associated with the senescence-high pixels without a previously well-established role in senescence, for example complement system factors (*C9, Cfh, Cfi, Hc*), kynurenine pathway related amino acid stress metabolism (*Gldc, Kyat3, Kynu*), and mitochondrial functional genes (*Sfxn1, Slc25a51*). Of these, several are involved in cell-cycle regulation, secretory phenotypes, stress response and immune-interaction (Figure 4e). Among the secreted proteins, *Fga, Fgb* and *Fgg* encode coagulation factors and are essential for the production of functional fibrinogen. A recent study using genome wide CRISPR/Cas9 screening identified coagulation factor IX (F9) as a key regulator of senescence[37]. Collectively, the data illustrate age-, sex-, and zone-specific senescence signatures.

**Figure 4.**
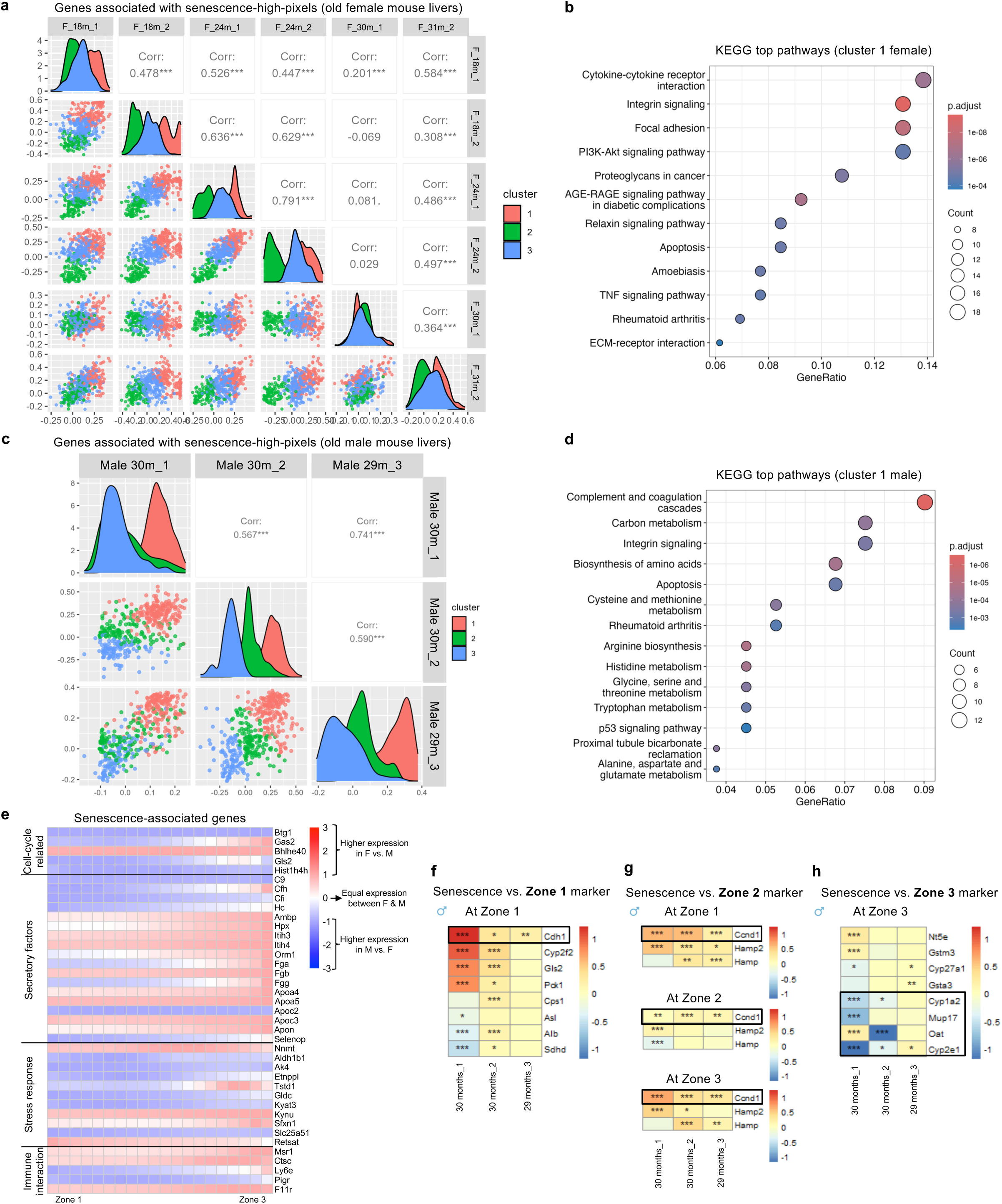
Significantly increased expression of senescence-associated genes identified across the liver lobule in old female and male mouse livers by Visium spatial transcriptomics. a. Replicative plots showing the correlation between each of the old WT female mice to assess reproducibility of detection of senescence-associated gene expression across biological replicates. Significant Pearson correlation coefficient (r) was calculated to quantify data consistency using cor.test() function in R, \*\*\**p* < 0.001. b. KEGG pathway enrichment analysis of senescence-associated genes in female mouse livers was performed using the enrichKEGG function in R. Pathways were ranked by enrichment *p* value. GeneRatio represents the proportion of input genes assigned to a given pathway. Count represents the number of input genes mapped to each pathway. Dot size corresponds to Count, and color indicates the adjusted *p* value (Benjamini-Hochberg correction). c. Same as a. except for male mice. d. Same as b. except for male mice. e. Heatmaps illustrating the expression pattern across liver lobule (x-axis) of up-regulated genes which are significantly associated with high-senescence-score pixels and without a previously well-established role in cellular senescence that were identified in old female (average expression of 18- and 24-month-old samples) and male (30-month-old samples) mice using zonation-aligned, cross-sample normalized data. These genes are categorized into 4 major functional groups associated with senescence (y-axis). Relative log_2_ fold-change was calculated for each gene at every zonation bin between sexes. A value of 0 (white box) indicates equal expression between sexes, positive values (red) indicate higher expression in females, and negative values (blue) indicate higher expression in males. f. Heatmap showing the correlation between high-senescence-score-pixels and the expression of zone 1 markers (y-axis) within zone 1 of each of the old WT male mouse livers (x-axis). Correlation coefficient ranging from −1.0 (prefect negative relationship, blue) to +1.0 (perfect positive relationship, red). T-distribution test: \**p* < 0.05, \*\**p* < 0.01, \*\*\**p* < 0.001. g. Heatmap showing the correlation between high-senescence-score-pixels and the expression of zone 2 markers (y-axis) across three liver zones of each of the WT male mouse livers (x-axis). Correlation coefficient ranging from −1.0 (prefect negative relationship, blue) to +1.0 (perfect positive relationship, red). T-distribution test: \**p* < 0.05, \*\**p* < 0.01, \*\*\**p* < 0.001. h. Heatmap showing the correlation between high-senescence-score-pixels and the expression of zone 3 markers (y-axis) within zone 3 of each of the WT male mouse livers (x-axis). Correlation coefficient ranging from −1.0 (prefect negative relationship, blue) to +1.0 (perfect positive relationship, red). T-distribution test: \**p* < 0.05, \*\**p* < 0.01, \*\*\**p* < 0.001.

We next asked the question: Does age-associated senescence correlate with age-related liver zonation changes? There was a significant positive association between senescence and zonation marker genes at zone 1 and zone 2, showing strongest correlation with *Cdh1* and *Ccnd1,* respectively, especially in males (Figure 4f, g; Extended Data Figure 4a). Interestingly, zone 2 marker *Ccnd1* also showed significant positive correlation with senescence at zone 1 and 3 in male (Figure 4g) and at zone 3 in female (Extended Data Figure 4b), which is consistent with the age-related expansion of zone 2 markers (Figure 1c-f). In contrast, an inverse association was observed between senescence and multiple zone 3 markers in both male and female livers (Figure 4h; Extended Data Figure 4c), suggesting profound and divergent roles of cellular senescence in age-associated zonation changes.

### Age-associated zonation disruption was recapitulated in *Ercc1^−/Δ^* mouse model

To determine if senescence plays a causal role in liver zonation changes, we queried whether livers from *Ercc1*^−/Δ^ mice (an established murine model of accelerated senescence of liver and other organs[33, 38]) recapitulated the zonation changes observed in aged WT mice. Bulk RNA-seq confirmed increased expression of SASP, inflammation, aging, and senescence-related genes in *Ercc1^−/Δ^* mouse livers relative to young WT animals (Figure 5a). Indeed, disrupted zonation patterning was shown by immunostaining in 4-month-old mutant mice (Figure 5b). Furthermore, GeoMx data revealed that, like aged WT mice (Figure 1b), *Ercc1*^−/Δ^ mouse livers displayed a large disruption of zone 2 gene expression (Figure 5c), in fact more so than that seen with aging. Consistent with the data in WT mice (Figure 4f, g), *Cdh1* and *Ccnd1* gene expression was significantly increased in *Ercc1^−/Δ^* livers (Figure 5d). Furthermore, an increased percentage of Cyclin D1 positive cells was observed in *Ercc1^−/Δ^* livers (Figure 5e). These data support our hypothesis that senescence promotes age-related changes in liver zonation, with concomitant effects on liver function.

**Figure 5.**
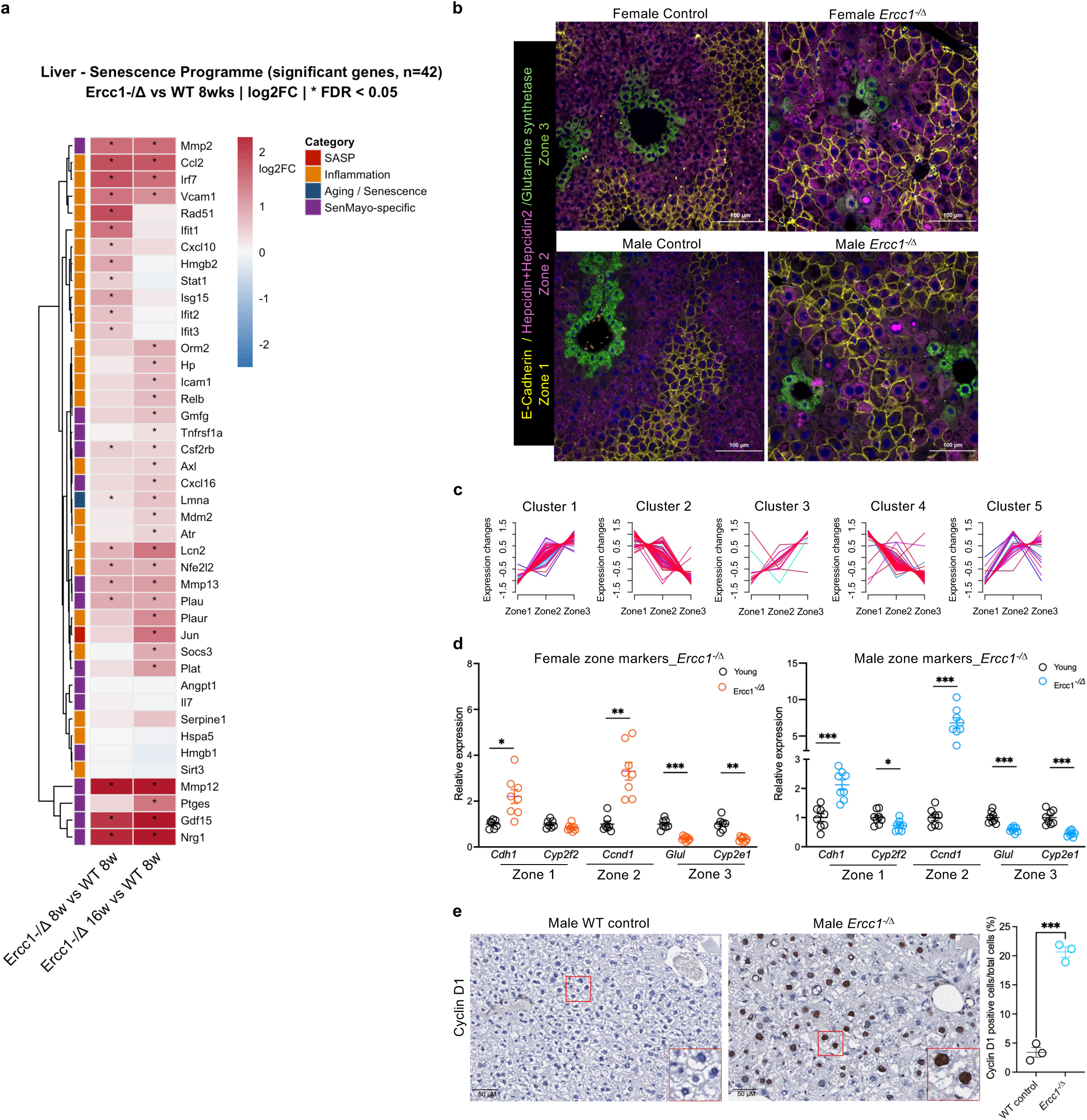
Age-associated zonation disruption was recapitulated in *Ercc1^−/Δ^* mouse model. a. Heatmap of differential expression of SASP, inflammation, aging/senescence and SenMayo genes (y-axis) identified in livers of *Ercc1^−/Δ^* (2 and 4-months-old; x-axis) relative to 2-month-old WT mice by bulk RNA-seq. n=3 mice per group. *FDR < 0.05. b. Immunofluorescent staining of zonation markers in mouse livers of *Ercc1^−/Δ^* mice at age of ∼ 4months and age-matched WT controls: E-Cadherin (yellow): periportal (zone 1), Hepcidin+Hepcidin2 (magenta): midlobular (zone 2), Glutamine synthetase (green): pericentral (zone 3). c. The same 215 DEGs across three liver zones identified using GeoMx digital spatial profiling in WT young livers (Figure 1b, top panel) were analyzed in *Ercc1^−/Δ^* mouse livers and their expression patterns aligned with the five gene clusters were illustrated. Gene expression patterns were most disrupted in zone 2 across all 5 clusters in *Ercc1^−/Δ^* mouse livers. Each line represents the expression variance of a single gene across the three zones. Warm colors (red and pink) indicate high confidence with which the gene belongs to that cluster, whereas cool colors (blue and green) indicate lower confidence. d. qPCR analysis of zone marker gene expression in young control and ∼ 4-month-old *Ercc1^−/Δ^* mouse livers. Each circle represents an individual mouse (n=7-8 per group). Left panel: female; right panel: male. Multiple unpaired *t*-tests and adjusted *p* values are presented as: \**p* < 0.05, \*\**p* < 0.01, \*\*\**p* < 0.001. Young WT controls were the same as used in Supplementary Figure 1e for female and Figure 1d for male. e. Representative images of immunohistochemical staining for the midlobular zone marker Cyclin D1 in male control and *Ercc1^−/Δ^* mouse livers. Two-tailed student *t*-test was applied for Cyclin D1 positive cell (%) quantification comparison, \*\*\**p* < 0.001. n=3 for WT young (∼4-month-old), n=3 for *Ercc1^−/Δ^* mouse (16 - 18 weeks) male mice. Young controls were the same as in Figure 1f.

### Zonation master regulator WNT signaling diminished in aged mouse livers

Upon identifying the impact of cellular senescence on age-related zonation changes, we next investigated the WNT signaling pathway, the master regulator of zonation, in aged mouse livers[22]. WNT signaling is well-established to be dominated by the interaction between endothelial cells and hepatocytes at pericentral zone (zone 3)[25]. This involves ligands of WNT2, WNT9b and RSPO3 secreted by endothelial cells that interact with receptors on hepatocytes through WNT/FZD/LRP coordinating with the RSPO3/LGR5/ZNRF3/RNF43 axis[21, 22, 25, 39].

In Visium data, *Wnt2* expression was decreased in livers of 18-month-old mice (Figure 6a). In addition*, Notum*, a negative regulator and direct target of the Wnt/β-catenin pathway[40], was significantly decreased in old WT mouse livers, particularly at pericentral zone 3 regions (Figure 6a). *Lect2* gene, which encodes hepatokine and immunomodulator LECT2, and a direct target gene of Wnt/β-catenin signaling in the liver[41], was also down-regulated at zone 3 (Figure 6a). This indicated diminished WNT signaling potentially contributing to the age-related zonation changes. This led us to further identity and characterize endothelial cells and hepatocytes in aged liver, the two key players of WNT signaling in dominating liver zonation.

**Figure 6.**
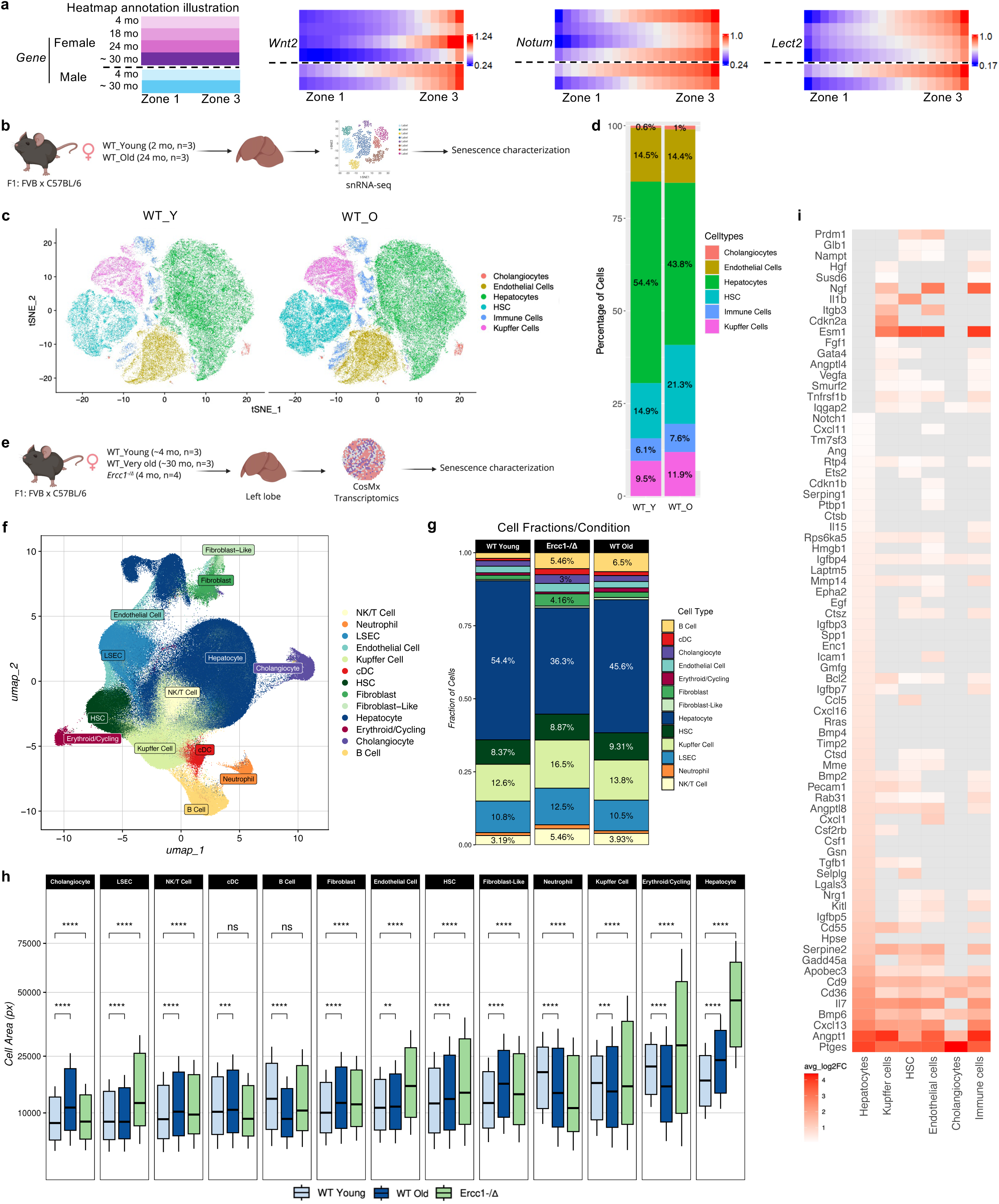
Identification of senescent cell types in aged mouse livers by snRNA-seq and CosMx spatial transcriptomics. a. Zonation master regulator WNT signaling diminished in aged mouse livers. Left: Illustration of how the three heatmaps of Visium transcriptomics data (to the right) are organized. Each row in the heatmaps represents a different age group of female mice (top 4 rows) and male mice (bottom 2 rows). On the x-axis, DEG expression is indicated across the liver lobule from zone 1 (left) to zone 3 (right). The three heatmaps illustrate age-related expression patterns of *Wnt2, Notum,* and *Lect2* across liver lobule. b. Schematic illustration of the experimental design of snRNA-sequencing. c. tSNE plot showing the major cell types detected in mouse livers by snRNA-seq. d. Stacked bar plot showing age-associated changes of hepatic cell composition detected by snRNA-seq. e. Schematic illustration of the experimental design of CosMx spatial transcriptomics. f. UMAP illustrating the identification of the major cell types in mouse livers detected by CosMx. g. Stacked bar plot showing age-associated changes of hepatic cell composition detected by CosMx. h. Cell area of different types of liver cells measured on CosMx images following cell segmentation (Seurat v5.0.1.9001). Px stands for pixels. One pixel is roughly 0.12 μM in diameter (or 0.0144 μM^2). *t*-tests were applied and *p* values are presented as: \*\**p* < 0.01, \*\*\**p* < 0.001, \*\*\*\**p* < 0.0001. i. Heatmap illustrating expression patterns of the gene panel used to identify senescence features (Supplementary Table 2) in liver cells by snRNA-seq. Grey bar represents the gene (y-axis) is not significantly differentially expressed for that specific cell type (x-axis).

Single-nucleus RNA-seq (snRNA-seq) (Figure 6b) and CosMx spatial transcriptomics (Figure 6e) were used to identify liver cells expressing senescence biomarkers among old and young samples. Notable changes in the cell type composition of aged livers relative to young were observed by snRNA-seq, with increased immune and hepatic stellate cells and decreased hepatocytes (Figure 6c, d; Extended Data Figure 5a). Similar change in cell composition between young and old WT mice was also observed by CosMx, as well as in the accelerated senescence *Ercc1*^−/Δ^ mice (Figure 6f, g; Extended Data Figure 5b). There was a significant enlargement of cell size across almost all cell types in old WT mice compared to young, which was more profound in *Ercc1*^−/Δ^ mouse liver cells, consistent with senescence-associated morphological changes (Figure 6h; Extended Data Figure 5c). Gene profiling of the snRNA-seq data revealed increased expression of numerous senescence-associated genes based on our 3-category senescence associated gene list (Supplementary Table 2) across multiple liver cell types, in particular hepatocytes, endothelial, hepatic stellate and Kupffer cells (Figure 6i).

### Senescent-like endothelial cells lose cell identity and marginally contribute to age-related zonation changes

Based on the zonation markers, we identified subclusters of zone 1+2 (periportal + midlobular) endothelial cells and zone 3+2 (pericentral + midlobular) endothelial cells (Extended Data Figure 6a, b). Using the 3-category of senescence-association genes strategy, we identified senescent-like cells in sub-zone endothelial cells (Extended Data Figure 6c, d). For zone 1+2 endothelial cells, we only detected *Itgb3, Prdm1* and *Gadd45a* significantly increased in both young and old senescent-like cells. For zone 3+2 endothelial cells, we only detected *Il7* significantly increased in old senescent-like cells, and an additional two senescence-associated genes (*Apobec3* and *Itgb3*) significantly increased in both senescent-like young and old endothelial cells.

Next, we turned to CosMx spatial transcriptomics for spatially better-defined endothelial cell population to investigate WNT signaling with age. After careful QC controlling for read depth variation and cell size (Extended Data Figure 7a), we focused our analysis on the pericentral region through identification of inner central vein hepatocytes and endothelial cells annotated by using hand-selected high confidence inner central vein (CV) regions (Extended Data Figure 7b). Hepatocytes and endothelial cells annotated in these inner CV regions grouped into well-separated subclusters that overlapped with expression of the pericentral zone 3 marker *Glul* (Extended Data Figure 7c).

Using a strategy similar to that used for identification of senescence in Visium and snRNA-seq data, we applied three categories of senescence-associated genes (SASP, cell cycle inhibitor and anti-apoptosis) to identify senescent cells, along with successful control of read depth variation related to cell size (Extended Data Figure 7d, e). We identified 66 senescent-like endothelial cells in the inner CV regions using CosMx in WT old samples (Extended Data Figure 7f). This small number of cells did not permit robust statistical analysis. Therefore, we did further analysis of these inner CV endothelial cells at the pseudo-bulk level. Inner CV ECs by CosMx had decreased expression of endothelial lineage marker genes such as *Cdh5, Kit and Tek/Tie2*, and endothelial functional genes such as *Fabp4, Kdr, Mrc1, Ephb4, Adgrf5* and *Lamp2*, suggesting age-associated loss of cell identity and function (Extended Data Figure 8a; Supplementary Table 4). Co-localization of WNT signaling genes (*Wnt7b, Wnt7a,* and *Wnt3*) (Note: WNT signaling key ligands *Wnt2* and *Wnt9b* were not included in the CosMx gene panel), and key WNT signal ligand *Rspo3* were reduced in *Cdh5*- and *Kit*-expressing endothelial cells in old WT mouse livers (Extended Data Figure 8b, c). These data suggest a potential role of senescent endothelial cells in contributing to age-related changes in zonation.

To test this more directly, *Ercc1* was deleted specifically in endothelial cells using *Tie2-Cre[42]*, to drive senescence selectively in these cells. *Tie2-Cre^+/−^;Ercc1*^−*/fl*^ mice showed increased expression of the zone 1 markers *Cdh1* (Extended Data Figure 8d), as well as zone 2 marker Cyclin D1 (Extended Data Figure 8e). These results indicated endothelial senescence due to *Ercc1* deletion led to disrupted zonation in this mouse model. Notably, *Tie2-Cre*^+/−^;*Ercc1*^−*/fl*^ mouse livers had less of an increase of the zone 2 marker Cyclin D1 compared to the increase in *Ercc1*^−*/Δ*^ mouse livers: 3.5% vs. 20.7% (Extended Data Figure 8e, Figure 5e), suggesting senescence in other cell types may contribute synergistically to *Ercc1*^−*/Δ*^ zonation disruptions.

### Senescent-like hepatocytes lose cell identity and contribute to age-related zonation changes

Since we only observed marginal impacts from pericentral endothelial cells on WNT signaling and zonation changes in aged liver, we further investigated the role of senescent hepatocytes. To establish senescence of hepatocytes beyond transcriptomics, primary hepatocytes were isolated from young and old WT and *Ercc1*^−/Δ^ mice. Hepatocytes from old and progeroid mice were enlarged, irregular in shape, with nuclear enlargement, features of senescent cells (Extended Data Figure 9a), with elevated expression of senescence, aging and inflammation-associated genes measured by bulk RNAseq (Extended Data Figure 9b) and qPCR (Extended Data Figure 9c). Lysotracker staining was increased in primary hepatocytes from old WT (Extended Data Figure 9d) and *Ercc1*^−/Δ^ mice (Extended Data Figure 9e) compared to young WT mouse cells, consistent with senescence-associated lysosomal biogenesis. *In vitro* culture of the primary hepatocytes significantly increased senescence marker gene expression (Extended Data Figure 9f).

To query whether hepatocytes residing in different zones of the liver have distinct senescence signatures, we clustered them based on zonation marker gene expression using the snRNA-seq data (Figure 7a; Extended Data Figure 9g). Notably, in young WT mice ∼11% of hepatocytes had undetectable levels of expression of the 11 zonation marker genes measured, which we labeled as no-zone hepatocyte-like population. This population expanded to ∼17% in aged WT livers (Figure 7b). This no-zone hepatocyte-like population showed significantly decreased HNF4α target gene expression, for example*, Ttr[43], Lipc[44], Cyp2c54[45] and Cyp3a11[46]*. The fraction of zone 2 hepatocytes also increased with age, while the fraction of zone 3 hepatocytes contracted (Figure 7b). RNA velocity analysis was used to predict cell fate changes that occur in hepatocytes with aging (Figure 7c). In young animals, zone 2 hepatocytes radiated towards other zones indicating that the zone 2 hepatocytes are the primary source of new hepatocytes across the liver lobule, consistent with published findings[26–28]. In old animals, there was evidence of zone 1 hepatocytes also contributing to zone 3 hepatocytes suggesting more need for regeneration in zone 3 and/or diminished capacity of zone 2 to fulfill that need. The no-zone hepatocyte-like population appeared to be primarily derived from zone 2 and, to a lesser extent, zone 1, particularly in old animals. The no-zone hepatocyte-like population had less RNA velocity than hepatocytes in other zones, suggesting these cells reside at an age-associated developmental end point with little capacity for regeneration. Gene expression profiling of the different hepatocyte clusters using the SenMayo gene panel revealed significantly increased expression of numerous genes in hepatocytes from old mice relative to young across all zone subclusters (Figure 7d). However, the strongest signal was observed in the no-zone hepatocyte-like population (young hepatocytes < old zone 3 < old zone 1 < old zone 2 << young no-zone << old no-zone).

**Figure 7.**
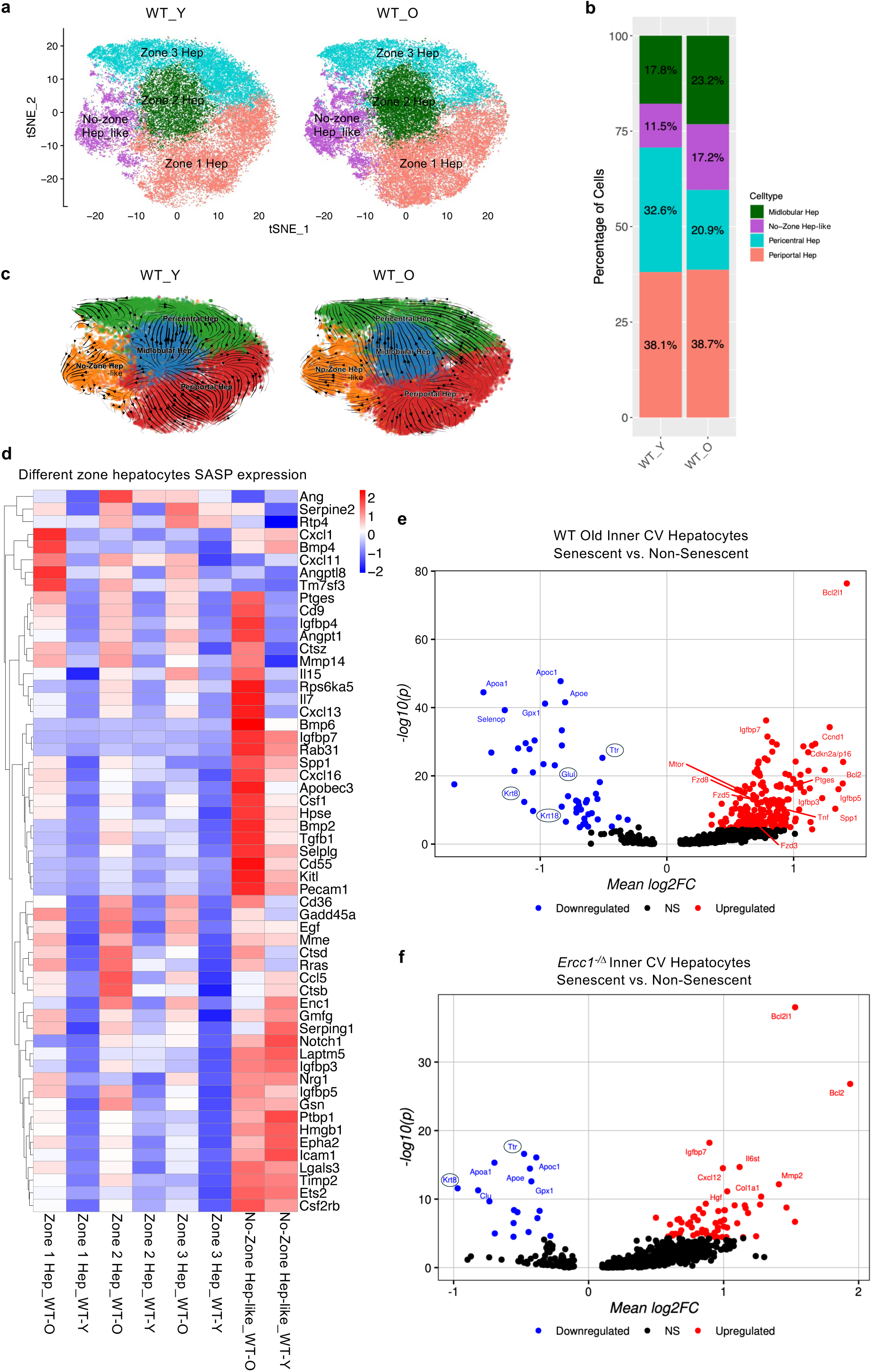
Characterization of senescent hepatocytes by zone from snRNA-seq and CosMx. a. tSNE plot from snRNA-seq data showing the identification of the hepatocyte clusters separated based on expression of zonation markers as described in Figure 6b. b. Stacked bar plot illustrating the abundance of hepatocytes representing each of the liver zones. c. RNA velocity analysis of young vs. old hepatocytes from snRNA-seq data. d. Heatmap of the expression of the SenMayo gene panel in hepatocyte subclusters from young vs. old WT mice across the different zones of liver. e. Volcano plot of DEGs in senescent vs. non-senescent hepatocytes identified using three senescence-categories strategy (e.g. SASP, cell cycle inhibitor and anti-apoptosis) within inner central vein (CV) regions of old WT mouse livers by CosMx studies. WT old n=3 mice. f. Volcano plot of DEGs in senescent vs. non-senescent hepatocytes identified using three senescence-categories strategy (e.g. SASP, cell cycle inhibitor and anti-apoptosis) within inner central vein (CV) regions of *Ercc1*^−/Δ^ mouse livers by CosMx studies. *Ercc1*^−/Δ^ n=4 mice.

Zone 3 appeared less affected in *Tie2-Cre^+/−^;Ercc1*^−*/fl*^ (*Glul* and *Cyp2e1*) (Extended Data Figure 8d) as in old WT and *Ercc1*^−/Δ^ mouse livers (Figures 1d, Figure 5d). The profound hepatocyte SASP profile (Figure 7d) led us to query if senescent zone 3 hepatocytes contributed to disrupted WNT signaling and zonation with aging. Senescent-like zone 3 hepatocytes identified in Inner CV regions by CosMx in old WT (Figure 7e, Extended Data Figure 7d-f, Supplementary Table 5) and *Ercc1*^−/Δ^ (Figure 7f, Extended Data Figure 7d-f, Supplementary Table 6) mouse livers showed significantly increased expression of *Cdkn2a, Bcl2, Igfbp7, Ptges and Spp1,* and loss of hepatocyte marker gene expression (*Glul, Ttr, Krt8, Krt18*), indicating loss of hepatocyte identity. This is consistent with findings that mice with hepatocyte-specific deletion of WNT signaling key factors LGR4/5 or RSPO blockade failed to express the zonation and pericentral metabolic genes including *Glul*, *Cyp2e1*[21]; and β-catenin knockout mice have down-regulated protein levels of GS (glutamine synthetase, encoded by *Glul* gene) and CYP2E1, and suboptimal liver regeneration[47]. β-catenin deletion also decreases expression of key hepatocytes transcription factors such as CCAAT-enhancer binding protein-α (C/EBPα) and hepatocyte nuclear factor-4α (HNF4α)[48]. In addition, knockout *Apc* gene to activate β-catenin signaling in mice leads to decreased expression of transthyretin (*Ttr*) and cytokeratin-18 (*Krt18*) expression, indicating repressed differentiation towards the hepatocyte lineage during development[49], highlighting the importance of balanced WNT signaling in liver development, homeostasis and regeneration. Although, there was a mixed pattern of expression of WNT signaling components (*e.g.*, genes were both up- and down-regulated), the net balance was diminished signaling (*e.g.,* significantly decreased co-localization of Wnt receptors and the hepatocyte marker *Glul*) in old WT mouse livers (Extended Data Figure 8f, Figure 7e). Collectively, these observations support a role for senescent zone 3 hepatocytes in age-associated liver zonation changes, closely tied to disrupted WNT signaling.

### Senescent hepatocyte signaling implicated in zone 2 marker expansion with age

We further investigate the role of senescence in zone 2 marker expansion in aged livers. Unsupervised cell clustering using a customized curated senescence gene list (Supplementary Table 2) revealed subpopulations of hepatocytes separated into distinct subclusters (Figure 8a). Expression mapping of three senescence-associated gene categories, e.g. SASP, cell cycle inhibitor and anti-apoptosis showed strong overlapping signals across all zones of hepatocytes, more pronounced in old samples (Figure 8b). Detailed analysis at the single gene level revealed senescence signatures in old hepatocytes across different liver zones (Figure 8c; Supplementary Table 7).

**Figure 8.**
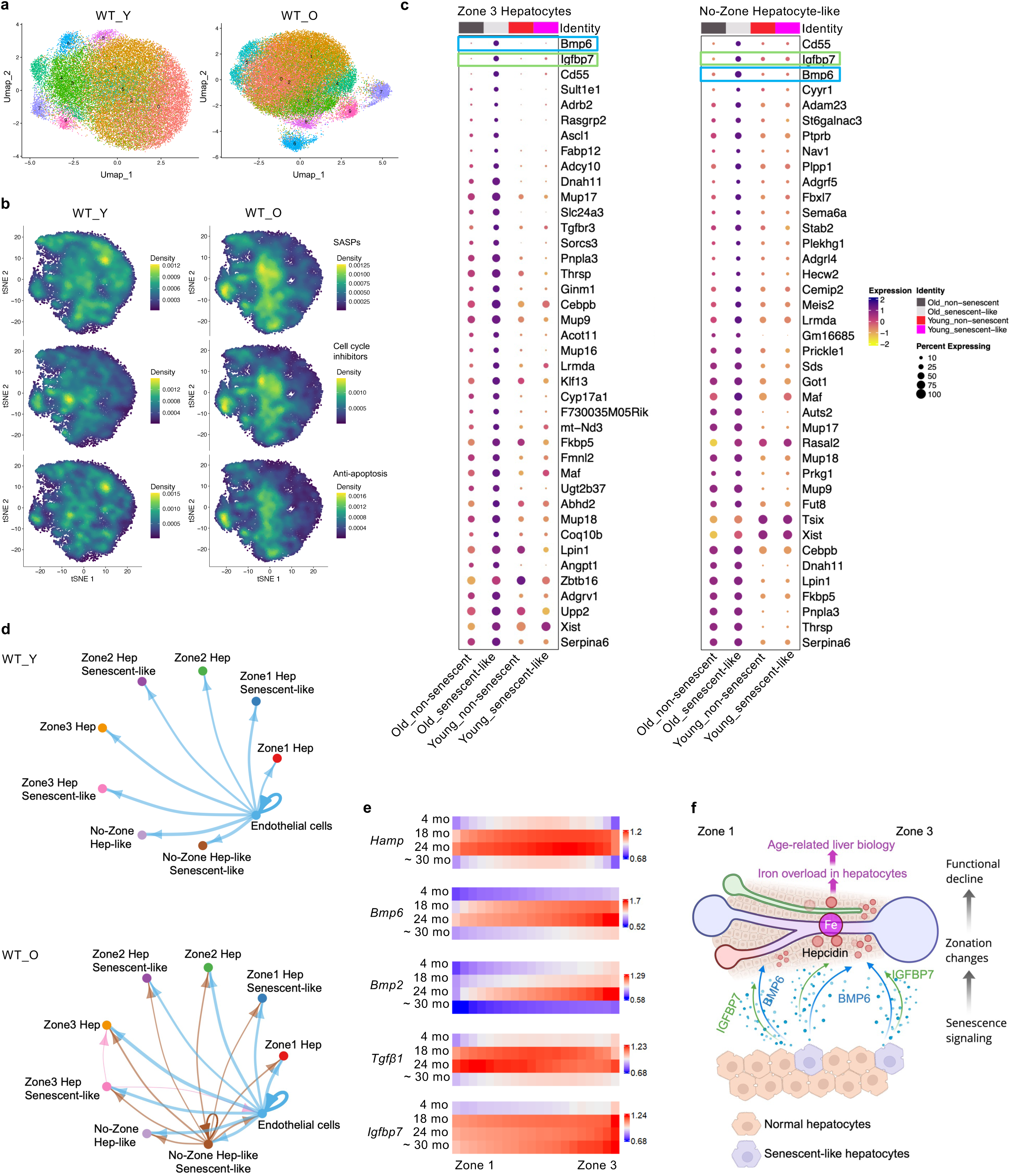
Senescent hepatocyte signaling implicated in zone 2 marker expansion with age. a. Unsupervised clustering of distinct subclusters of hepatocytes from young vs. old WT mice defined by differential expression of senescence-associated genes (Supplementary Table 2). b. Density plots of hepatocytes from young (left) and old (right) WT mice illustrating cells with highest expression of three senescence-associated gene classes (SASP, cell cycle inhibitors, anti-apoptosis genes, from top to bottom). c. Dot plots illustrating the expression of the top senescence-associated genes in old senescent-like vs. non-senescent hepatocytes in zone-3 hepatocytes (left) and no-zone hepatocyte-like population (right). d. Circle plots illustrating the BMP signaling pathway communication pattern between endothelial cells and senescent-like vs. non-senescent hepatocytes from different zones of the liver from young (top) vs. old (bottom) WT mice. e. Heatmap illustrating the age-related expression patterns of the listed genes (on the left) across liver zones. Each row in the heatmap represents the expression pattern of the gene across liver lobule from zone 1 to zone 3 (x-axis) in different age group (y-axis) of female mice by Visium transcriptomics. f. Schematic illustration of the effects of BMP6 and IGFBP7 secreted by senescent-like hepatocytes on increasing hepcidin expression and iron deposition in aged mouse livers.

Bone morphogenetic protein 6 (*Bmp6*) was the SASP factor most strongly upregulated with age in no-zone hepatocyte-like cells (Figure 7d, Figure 8c). The BMP signaling pathway is one of the top pathways identified in old livers using CellChat analysis (data not shown). BMP6 is a central transcription factor that through SMAD signaling maintains iron homeostasis by regulating hepcidin. Hepcidin, encoded by Zone 2 marker gene *Hamp*, is the master regulator for systemic iron balance through degradation of the iron exporter ferroportin[50]. Endothelial cells are the dominant source of BMP6 in the liver[51] and CellChat analysis of our snRNA-seq data in young mouse livers recapitulated this finding showing BMP signaling originated from endothelial cells signaling hepatocytes (Figure 8d). However, in old female livers, a strong signal also emerged from senescent-like no-zone hepatocyte-like cells targeting all the rest of hepatocytes. A subtle but detectable BMP signal was also observed stemming from senescent-like zone 3 hepatocytes signaling to non-senescent zone 3 hepatocytes and endothelial cells in old samples (Figure 8d). This snRNA-seq result is consistent with what we observed in Visium spatial studies, in which zone 2 marker *Hamp* was increased with age with concomitant upregulation of *Bmp6*, *Bmp2*, especially at zone 2 and zone 3 regions (Figure 8e).

Increased *Tgfβ1* expression was observed as well (Figure 8e), which also stimulates hepcidin expression and functions in iron metabolism regulation[52]. IGFBP7 is a key SASP component and can induce cellular senescence through regulating insulin, IGF, and Activin A as well its downstream SMAD pathways[53]. High levels of IGFBP7 are detected in human non-alcoholic fatty liver disease (NAFLD) and liver fibrosis patients[54]. IGFBP7 promotes liver disease progression by regulating ferroptosis; hence, depletion of IGFBP7 significantly reduces iron deposition, lipid peroxidation and ferroptosis in zebrafish[54]. *Igfbp7* expression was significantly increased in senescent-like hepatocytes across all hepatic zones (Figure 8c and e; Supplementary Table 7) and also in Inner CV senescent hepatocytes in both WT old and *Ercc1^−/Δ^*livers at zone 3 by CosMx (Figure 7e, f) (Note: *Bmp6* was not in the CosMx gene panel). We propose that increased BMP6 and IGFBP7 by senescent-like hepatocytes promotes hepcidin expression, leading to decreased iron export out of hepatocytes and increased iron deposition inside of hepatocytes in aged livers, especially at zone 3 hepatic regions, leading to age-related liver function decline (Figure 8f).

## Discussion

Senescent cells are strongly implicated in the pathophysiology of numerous age-related chronic diseases including metabolic dysfunction-associated steatosis, inflammation, fibrosis and HCC[7]. However, their functional impact on aging biology, as opposed to pathophysiology, is less well understood. In this study, single nucleus RNA-seq and spatial transcriptomics were applied to identify and characterize senescent cells in mouse livers across lifespan to interrogate their mechanistic role in driving liver aging biology.

In aged liver, we identified cell composition changes and cellular hypertrophy consistent with previous reports[34]. Hepatocytes, endothelial, hepatic stellate and Kupffer cells acquired senescence features with normal aging, also consistent with previous findings[4, 5, 55]. We further identified that senescence signatures are age-, zone- and sex-specific. Senescence appeared in middle-age, increased over the next quintile of life and titered down in the last quintile of life. This indicates that the third quintile of life might be the appropriate time to administer senotherapeutics to prevent changes associated with liver aging. The reduction in senescence signal in the oldest animals could be due to the ‘centenarian’ effect: late-life survivors having attenuated aging biology and chronic disease.

With aging, senescent cells appear across the entire liver lobule and formed clusters especially at advanced age (Figure 3d and h; Extended Data Figure 3b and f) consistent with senescent cells corrupting their local tissue microenvironment and spreading senescence. The proportion of senescent cells seen in the murine liver with aging (<10%) is small, consistent with senescent cells exerting their effects by secreted SASP factors that are pro-inflammatory and tissue disrupting[8]. With aging, a population of hepatocytes with no clear zonation identity expanded and these cells are the most proinflammatory.

With aging, liver zonation is known to be disrupted. In murine liver, we found that zonation changes are detected by middle-age (18 months) including increased expression of zone 2 markers *Hamp* and *Ccnd1* but diminished expression of zone 3 markers such as *Glul* and *Cyp2e1* (Figure 1; Extended Data Figure 1). This coincides with increased expression of senescence genes, including *Cdkn2a* as we and others report[5]. The co-occurrence of senescence and loss of zonation suggests a potential mechanistic link. Indeed, we found senescence signatures were closely spatially related to zonation changes within the liver, with a positive correlation with zone 1 and zone 2 markers but an inverse correlation with zone 3 markers (Figure 4f-h). Zonation disruption is also coincident with senescence in *Ercc1*^−/Δ^ mice, a model of human progeria and accelerated liver senescence[38, 56], supporting a causal role of senescence in driving loss of zonation (Figure 5).

With aging, zone 3 is particularly affected, likely due to its role in xenobiotic and detoxification. Zone 2 serves as a regenerative reservoir to replenish ‘retired’ hepatocytes [26–28]. In our studies, zone 2 hepatocytes, the primary source of maintaining liver homeostasis, retained proliferative capacity as suggested by RNA velocity results (Figure 7c), but became more heterogenous and inflammatory with age (Figure 7d; Figure 8b). These cells do not completely lose their zone 2 signature nor do they gain the full zone 3 or zone 1 signatures, or functional profile necessary to fulfill the zone’s canonical functional demands. This may explain why with aging, migration of zone 2 hepatocytes to zone 1 and 3 needed for regeneration is less efficient (Figure 7c). This also explains the age-related expansion of zone 2 with contraction of zone 3 (Figure 1c, d). These data indicate that age-related spontaneous cellular senescence in the liver drives loss of hepatocyte identity and function, contributing to maladaptive regeneration and compromised metabolic resilience.

WNT signaling, the master regulator of liver zonation, is dominantly regulated by endothelial cells and hepatocytes at pericentral regions (zone 3). Deletion of *Ercc1* specifically in endothelial cells to drive senescence in these cells[42] indeed drives age-related changes in zone 2 marker expression (Extended Data Figure 8e), as expected. However, the impact on zonation patterns in these mice was rather marginal, as no significant changes were observed in zone 3 regions (Extended Data Figure 8d). In systemic *Ercc1^−/Δ^*mutant mice, zonation changes mimic those of aged livers in both zone 2 and zone 3 (Figure 5d; Figure 1d). This suggests that senescent hepatocytes, characterized herein, contribute to age-related changes in zone 3. This is consistent with disrupted WNT signaling factors, providing a mechanism of impaired hepatocyte function (Figure 6a). Studies identified zone 2 hepatocytes as the key player in maintaining liver homeostasis through (IGFBP2-mTOR-CCND1) axis[27]. Elevated expression of *Igfbp2* and *Ccnd1* in senescent hepatocytes could potentially drive altered zonation in the aged mouse livers through mTOR signal, especially in male mice.

Liver stores the highest amount of iron in the body which makes it vulnerable to iron toxicity. Age-related iron overload has been increasingly recognized as a public health concern[57]. Iron levels increase in old age, and old female mice have as much as 2-3 fold more of iron in their livers than males, especially in certain mouse strains[58]. Age-related iron accumulation shifts metabolic patterns revealed by metabolomics analysis and resets circadian clock[59]. Aging is a risk factor for iron-induced liver injury including inflammatory cell infiltration, cell death, extracellular matrix deposition and sinusoidal defenestration[60]. Higher iron levels are present in patients with nonalcoholic fatty liver diseases and other chronic liver diseases[61]. BMP signaling plays a key role in regulating hepatic iron metabolism by promoting the expression of the hormone hepcidin (encoded by *Hamp* gene, a zone 2 marker)[31]. *Bmp6* expression was significantly increased in senescent hepatocytes, which aligns with the age-related zone 2 marker expansion identified using multiple methods in our study (Figure 1 and 8). Ferroportin, the only known iron exporter in mammals[57], is downregulated by hepcidin through triggering its internalization and degradation[31, 50], therefore it is highly likely that increased hepcidin stops iron export and traps iron inside the hepatocytes leading to iron overload in aged liver. In addition, increased *Igfbp7* by senescent hepatocytes (Figure 7 and 8) exacerbates iron deposition[54], which combines with reactive oxygen species resulting in increased hydroxyl radicals, lipid peroxidation, DNA and protein damage, leading to liver dysfunction[61].

In summary, our data provide potential causal mechanistic links between cellular senescence and disrupted liver zonation associated with aging. Furthermore, the data illuminate several opportunities for intervention to restore age-related decline in liver regeneration and function including senotherapeutics, mTOR inhibitors and iron chelators.

### Limitations

We mainly used transcriptomics data in this study. Many key components of the signaling pathways addressed in this study are regulated post-translationally. In this study, we mainly focused on pericentral (zone 3) senescence in endothelial cells and hepatocytes and their potential role in WNT signaling regulation and zonation changes due to limitations of the study platforms. However, we also observed significant changes in other important signals such as Notch, which has been reported to play critical role in liver zonation and regeneration[23], therefore warrants further investigation in our studies. The findings in this study need to be tested in human liver samples.

## Materials and Methods

### Mice

All mouse handling and mouse tissue collection were carried out in compliance with regulations and IACUC-approved protocols at the University of Minnesota. Experimental mice were generated by breeding inbred C57BL/6J with inbred FVB/n mice resulting in progeny that are F1 hybrid and genetically identical. Mice were housed in the specific pathogen-free barrier facility at University of Minnesota until they reached the desired age for tissue collection. Mice were euthanized using CO_2_ and the liver tissue was excised and processed immediately either subjected to snap-frozen in liquid nitrogen, or the left lobe was fixed with 10% formalin for 24 hours and stored in 70% Ethanol at 4°C until paraffin embedding.

### GeoMx Digital Spatial Profiler Studies

Formalin-fixed paraffin-embedded mouse liver tissues were sectioned at 5 μm and mounted on positively charged 25 x 75 mm glass slides. Liver sections across 4 slides were processed simultaneously within 2 days of sectioning. The GeoMx Mouse Whole Transcriptome Atlas (NanoString 121401103) was used to generate libraries. Slide processing was performed per the manufacturer’s instructions (NanoString document MAN-10150-4) with the indicated notes and exceptions. Detailed experimental procedure and data analysis strategy are listed in Supplementary Methods.

### Visium Transcriptomics Studies

Distinct procedures were used for Visium direct mount (*n*=9 mouse livers) and CytAssist (*n*=4 mouse livers) methods. Both workflows followed the instructions of their manufacturers (10x Genomics documents CG000409 Rev. D and CG000407 Rev. E for direct mount; documents CD000520 Rev. C and CG000495 Rev. F for CytAssist). Detailed experimental procedures are in Supplementary Methods.

### Visium Data Analysis

After superpixel imputation (∼5,000 genes per sample) by iStar[62], we performed a down sampling step to reduce computational burden while retaining biologically relevant markers. We constructed a 1,000-gene matrix by preserving all mandatory genes, including housekeeping genes, immune markers, endothelial and fibroblast signatures, zonation markers, and senescence-related genes (SASP, cell-cycle inhibitors, and anti-apoptosis genes). The remaining genes were included as supplemental features primarily for PCA-based artifact estimation. Each Visium tissue slide was then manually cropped to retain only high-quality tissue regions for downstream analysis.

Following cropping, we applied preprocessing steps to clean data of outliers and artifacts. First, we performed per-gene clipping independently for each sample by thresholding expression values at the 5th and 98th percentiles to limit the effect of extreme values. We then computed total pixel intensity and applied intensity-based masking to flag outlier pixels outside the central distribution, which were optionally excluded in later steps.

In a subset of samples, we applied an additional endothelial-based masking step to identify pixels corresponding to large blood vessels. Using endothelial marker genes (*Cdh5, Vwf, Vcam1, Cd31, Cd144, Flt4,* and *Hba-a2*), we computed an endothelial expression score per pixel and modeled its expected value as a function of total intensity. Pixels with high residual endothelial signal (top 1%) were annotated as blood vessel pixels and masked for downstream analyses.

We then normalized pixel-level expression using a square-root variance-stabilizing transformation followed by housekeeping normalization. To further reduce technical variation, we performed PCA on housekeeping genes and identified PC1 as the dominant artifact axis. For each gene, we regressed expression on PC1 and removed the fitted technical component, preserving biological signal while reducing technical noise.

To characterize spatial organization, we computed a per-pixel zonation score using periportal and pericentral marker genes, forming a continuous pseudotime from zone 3 to zone 1. Scores were quantile-normalized across samples to ensure a consistent 0–1 scale, and pixels were binned into balanced zonation groups for downstream analysis.

For differential expression, we generated pseudobulk profiles by averaging expression within each sample and zonation-defined region (periportal, midlobular, pericentral). Samples were grouped by age (<12 months, 12–24 months, >24 months), and differential expression was performed using limma[63], modeling expression as a function of age and genotype. This framework allowed us to identify zone-specific age-associated transcriptional changes.

To quantify senescence, we computed per-pixel scores based on SASP, cell-cycle inhibitor, and anti-apoptosis gene signatures. In addition, *Mki67* filtering was applied to mask out pixels with *Mki67* expression level in the top 95^th^ percentile. Scores were converted to percentile ranks using pooled wild-type reference distributions and combined into a weighted senescence score. Pixels with high senescence scores were mapped spatially and correlated with gene expression to identify senescence-associated transcriptional programs. Detailed data analysis strategy is listed in Supplementary Methods.

### snRNA-seq Studies

Single nuclei were isolated from snap-frozen mouse liver tissues following the 10X Genomics protocol: CG000505_Chromium_Nuclei_Isolation_Kit_UG_RevA.

### snRNA-seq Data Processing and Analysis

The 10x Genomics FASTQ files were processed using Cell Ranger (v9.0) with the following parameters: --localcores=24, --localmem=200, --create-bam=true, --chemistry=SC3Pv3, and -- expect-cells=20000. Seurat (version 5.0) was used to convert 10x counts to a Seurat object. Preprocessing was performed to extract high-quality cells by assessing feature RNA counts and mitochondrial content. Cells with nFeature_RNA counts >1000 and <4000 and mitochondrial content <1% were selected for downstream analysis. Counts were normalized using NormalizeData with default parameters and 2000 variable features identified using the FindVariableFeatures function, then used for integration purposes. One of the samples from an old mouse was used as a reference to integrate data. Thirty principal components were calculated using the RunPCA function and Umap coordinates were calculated using the first 15 principal components using the RunUMAPfunction[64]. Neighbors were calculated on the first 15 principal components as well, using the FindNeighbors function. The FindClusters function was applied to calculate clusters at a resolution of 0.2. Harmony[65] available at https://github.com/immunogenomics/harmony was used to calculate harmony components and based on the first 15 components, a t-SNE map was created using the RunTSNE function in the Seurat package. Detailed data analysis strategy is listed in Supplementary Methods.

### CosMx Spatial Molecular Imager Studies

Formalin-fixed paraffin-embedded mouse liver tissues were sectioned at 5 μm thickness and mounted on Leica BOND PLUS glass slides. Five liver samples were mounted on each of 2 slides and both processed simultaneously within 3 days of sectioning. Slide processing was performed as instructed by the manufacturer (NanoString document MAN-10184-2). Detailed experimental procedure is listed in Supplementary Methods.

### CosMx Data Analysis and Statistics

Data from the two CosMx slides were analyzed in Seurat v5.0.1.9001. Log-normalization and SCTransform normalization were performed at the level of individual tissue samples. SCTransform included cell size as a variable to regress to help control for read depth variation across differentially sized cells. Cells with <100 reads and 50 distinct transcripts were filtered out prior to integration with Harmony using 30 principal components. Unsupervised clusters were generated using dimensions 1:15 for FindNeighbors and RunUMAP, and a resolution=1 for FindClusters. Each unsupervised clustering was annotated manually with cell-specific marker genes and annotations were subsequently confirmed using the top 20 most cluster-specific marker genes with CellKB. Inner CV (inner central vein) regions hepatocytes and endothelial cells were defined using a small set of hand-selected high confidence inner CV regions. Hepatocyte and endothelial cell annotated cells in these regions grouped to specific unsupervised clusters, which demonstrated tight spatial clustering. Thus, all cells in these clusters were re-labelled as inner CV cells. Differential expression was quantified with Seurat’s FindMarkers function on the Wilcox setting. Colocalization was tabulated using a scaled ratio of the Mander’s Overlap Coefficient (MOC) of each gene pair against a distribution of MOCs from pairs of genes with similar expression serving as a background calibrator. The complete scripts for this are available at: https://github.com/katlande/scCoExpress.

### Statistical Analysis

Statistical analyses were performed using GraphPad Prism11. Mean ± SEM was presented. For two groups comparisons, two-tailed student *t*-test was applied. For multiple unpaired *t*-test, correction for multiple comparisons using the Holm-šídák method and adjusted *p* values were presented. For more than two groups comparisons, one-way ANOVA was used and corrected by Dunnett’s test or Tukey’s test. Exact sample sizes, repeating times, error bars and statistical methods are provided in detail in the related Figure legends and Extended Data Figure legends.

## Data availability

All RNA-seq and Spatial transcriptomics datasets generated in this study have been deposited in the SenNet consortium data repository: https://data.sennetconsortium.org/. All the datasets and sample IDs are listed in Supplementary Table 8.

## Code Availability

Visium data analysis pipeline: https://github.com/alixha-rx/Liver_Project

## Author Contributions

L.L., L.J.N, and P.D.R. conceived and designed the study. L.L., S.P., J.H., L.A., M.K., J.B., and A.L. carried out the experiments. L.L., A.H., and P.A. contributed to senescence-associated reference gene list. M.J.S. and C.M.C. designed the custom CosMx codeset. L.L., N.Z., A.A., S.A., K.L., M.L., K.G.E., A.H., Z.M., J.Z., S.P., N.P., M.E.B., and O.A. carried out the data analysis. L.L., N.Z., A.A., S.A., K.L., M.L., K.G.E., A.H., Z.M., X.D., J.W., P.D.R., and L.J.N. contributed to data interpretation. L.L. wrote the manuscript, and L.J.N. reviewed and edited the manuscript with feedback from all the authors. All authors reviewed and approved the final manuscript.

## Acknowledgement

We are very thankful for all the support from the University of Minnesota Masonic Institute on the Biology of Aging and Metabolism (MIBAM) mouse core members Ryan O’Kelly, Sara Mcgowan, MacKenzie Beer and Alan Mangen; the technical support from Tra Kieu, Jake Kircher, Joshua Carey and Karah Shirley in the Niedernhofer lab; data analysis support from Shamsed Mahmud. This work was supported by the resources and staff at the University of Minnesota University Imaging Centers (UIC, RRID: SCR_020997), University of Minnesota Genomics Center (UMGC, RRID: SCR_012413), and the core members Pat Grady, Fernanda Rodriguez, Colleen Forster, Laurie Reichel, Scott Horsfall, Erin Hudson, Anna Peyla, Steve Schnell, Patrick Willey and Grant Barthel. We appreciate the insightful discussion with Drs. Douglas Mashek, Andrew Nelson, Byoung Uk Park, Lei Zhang, Christina Camell, and In Hwa Jang. We are thankful for the support from Dr. Davis Seelig, Katalin J. Kovacs, and Paula Overn at Comparative Pathology Shared Resource at University of Minnesota. Illustrations were generated using BioRender.

---

**Extended Data Figure 1.**
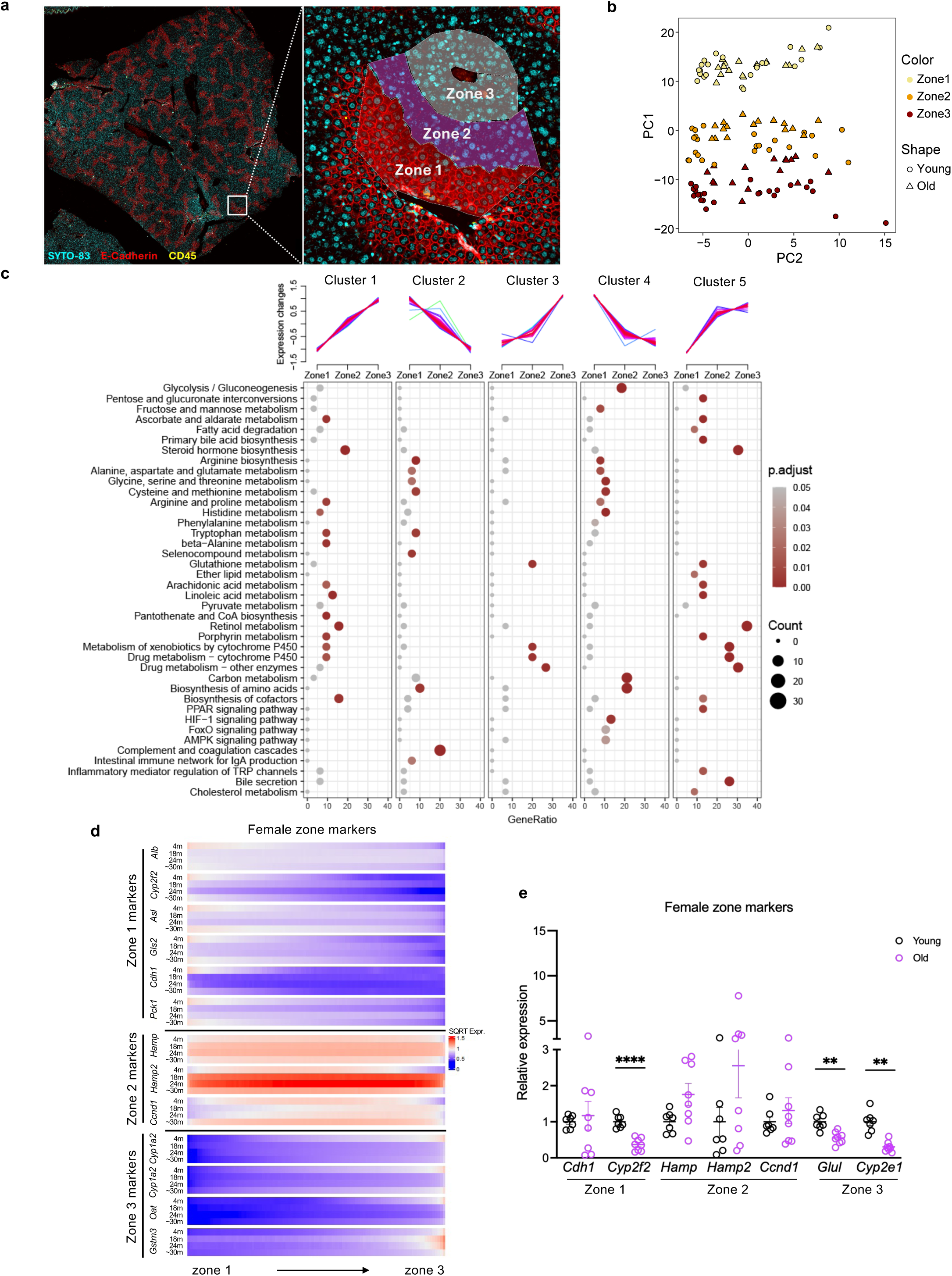
Age-related zonation changes identified by multiple methods (related to Figure 1) a. Representative image demonstrating ROI selection strategy utilized in GeoMx transcriptomics in liver samples immunostained for E-Cadherin (in red) to define periportal (zone 1) regions. This strategy combined with anatomical structures was used to manually define zones 1, 2 and 3. Left panel white box is one set of ROIs demarcating three zones, illustrated in higher magnification in the right panel. b. PCA plot showing clustering of the different hepatic zones across the GeoMx ROIs selected from young (circles) and old (triangles) WT mouse livers. c. Five gene clusters were identified from GeoMx digital spatial profiler based on the expression patterns of the 215 DEGs across the three hepatic zones in WT young livers (same as shown in Figure 1b). For example, cluster 3 is defined by gene expression greatest in zone 3 with roughly equivalent, and much lower, gene expression in zones 1 and 2. Each line represents the expression variance of a single gene across the three zones. Warm colors (red and pink) indicate high confidence with which the gene belongs to that cluster, whereas cool colors (blue and green) indicate lower confidence. The dot plots below each cluster show the results of pathway enrichment analysis based on the genes assigned to the corresponding cluster. KEGG pathway analysis of each gene cluster identified significantly enriched pathways that differed across the various clusters. The dot size represents the number of genes enriched in each pathway. The gene ratio (x-axis) indicates the proportion of genes enriched in a given pathway relative to the total number of genes within the cluster. The dot color represents the statistical significance (*p* value) of the enrichment (the darker the red the greater the significance). Hypergeometric test with adjusted *p* value < 0.05 defined as significant. d. Heatmaps showing the gene expression pattern of zone markers (y-axis) across the 3 liver zones (x-axis) in 4-, 18-, 24-, and ∼30-month-old female mouse livers (indicated by row) measured by Visium transcriptomics studies. e. qPCR analysis of zone marker gene expression in young and old female mouse livers. Each circle represents an individual mouse (n=7-8 per age group). Multiple unpaired *t*-tests and adjusted *p* values are presented as: \*\**p* < 0.01, \*\*\*\**p* < 0.0001.

**Extended Data Figure 2.**
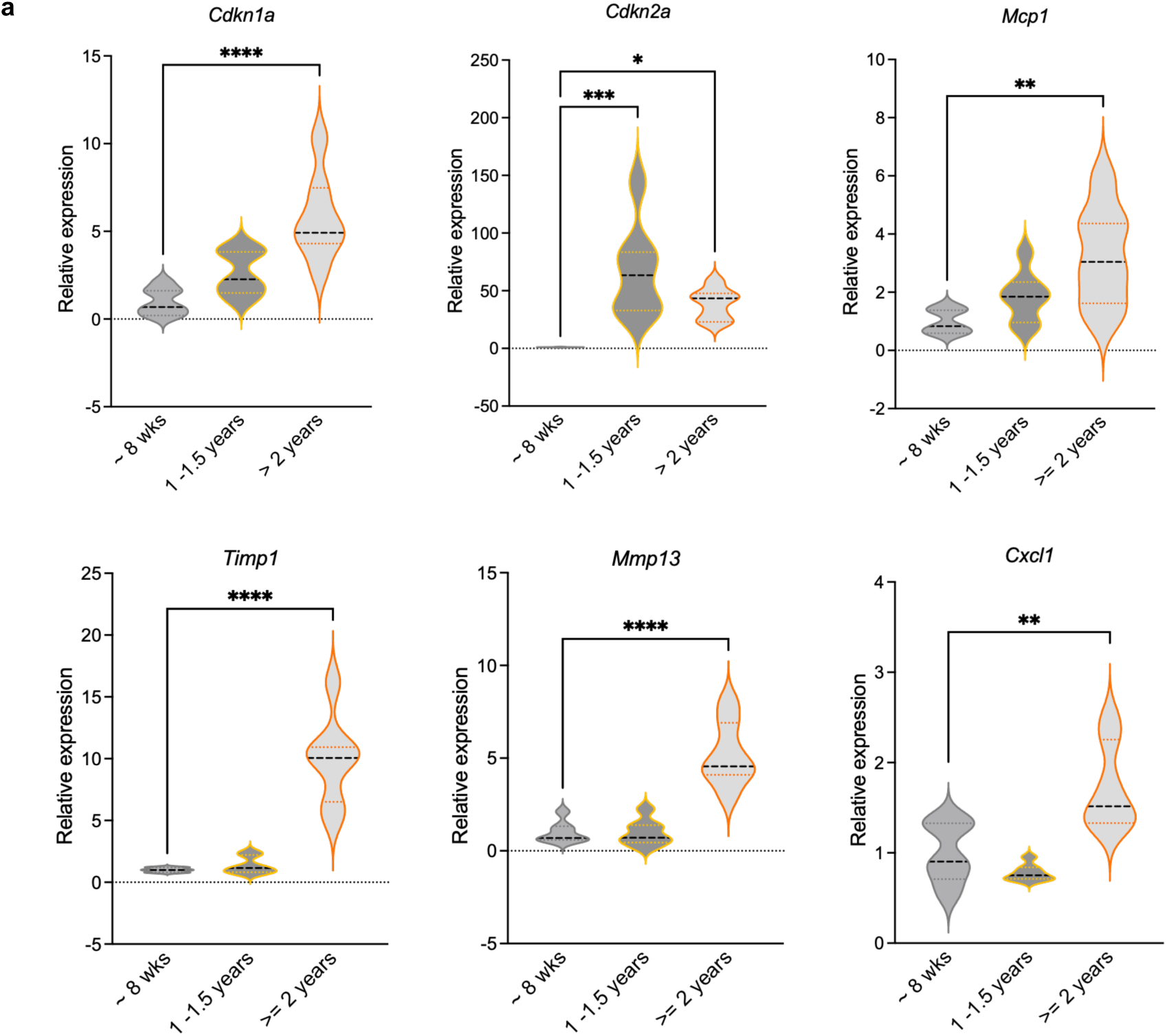
Expression of senescence-associated genes was significantly increased in livers of old mice relative to young (related to Figure 2) a. Increased expression of senescence-associated genes in aged WT mouse livers relative to young mouse livers measured by qPCR. n=7 per age group. One-way ANOVA analysis: \**p* < 0.05, \*\**p* < 0.01, \*\*\**p* < 0.001, \*\*\*\**p* < 0.0001.

**Extended Data Figure 3.**
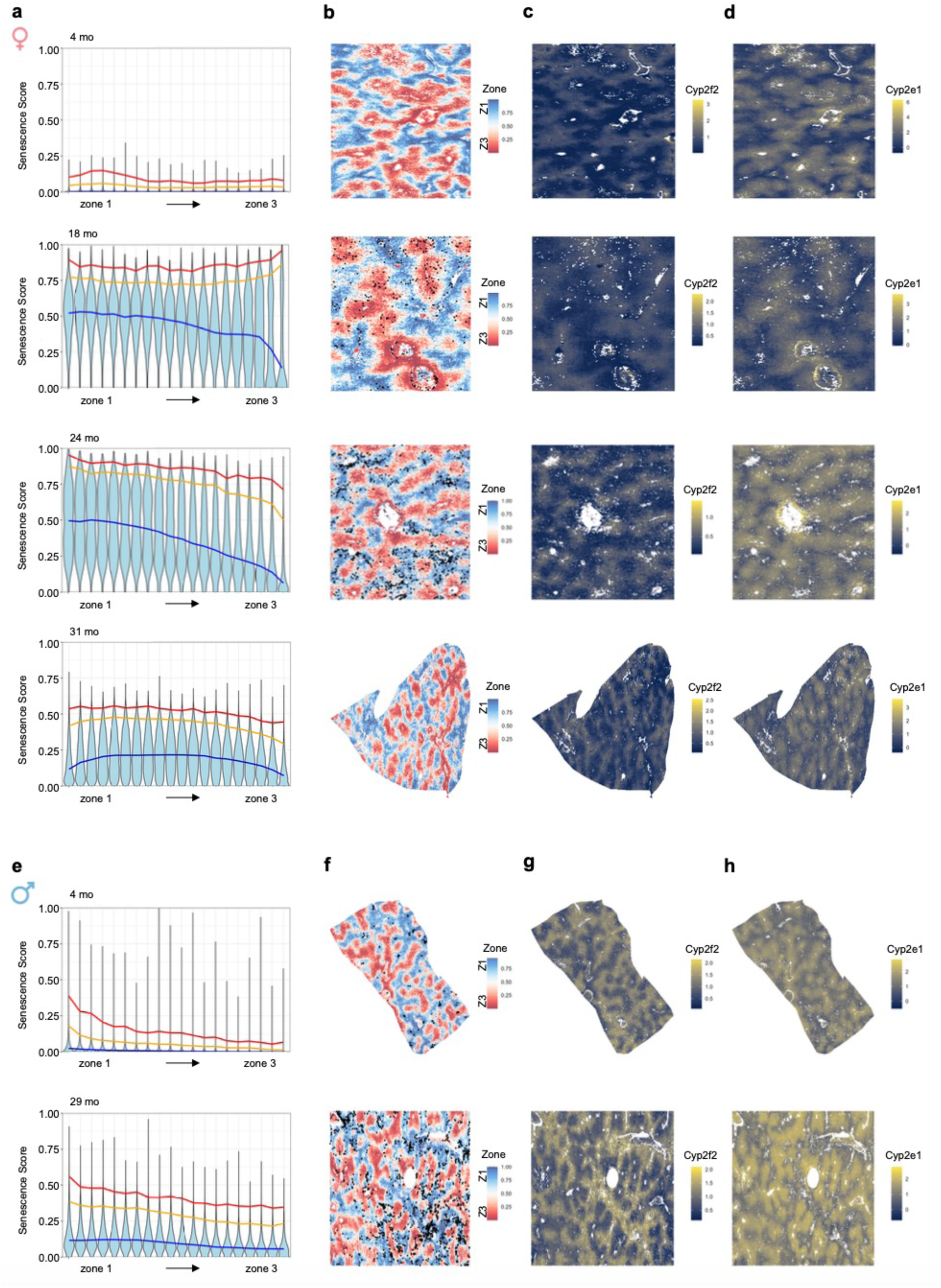
Age-, sex- and zone-related senescence signatures identification by Visium spatial transcriptomics (related to Figure 3) a. Violin plots showing senescence scores across the liver lobule from zone 1 to zone 3 (x-axis) in different age groups (4, 18, 24 and 31-months old, by panel top to bottom) of WT female mice. Liver zones were divided into 20 bins from zone 1 to zone 3, and each violin line represents the distribution of senescence score for each zonation bin. Blue line represents 50^th^ percentile, orange: 95^th^ percentile, and red: 99^th^ percentile. b. Spatial mapping of senescence-high pixels (black dots represent the top 98^th^ percentile of senescence score pixels in 24-month-old samples at a cutoff threshold of 0.86 [in Figure 3a]) in the zones of WT female mouse livers. Zone 1 blue, zone 3 red, at 4, 18, 24 and 31-months of age (top to bottom). c. Spatial distribution of gene expression of the zone 1 marker *Cyp2f2* in WT female mouse livers at 4, 18, 24 and 31-months of age (top to bottom). d. Spatial distribution of gene expression of the zone 3 marker *Cyp2e1* in WT female mouse livers at 4, 18, 24 and 31-months of age (top to bottom). e. Same as a, but for WT male mice at 4 and 29-months of age. f. Same as b, but for WT male mice at 4 and 29-months of age. (Black dots represent the top 98^th^ percentile of senescence score pixels in 30-month-old samples at a cutoff threshold of 0.41 [in Figure 3b]). g. Same as c, but for WT male mice at 4 and 29-months of age. h. Same as d, but for WT male mice at 4 and 29-months of age.

**Extended Data Figure 4.**
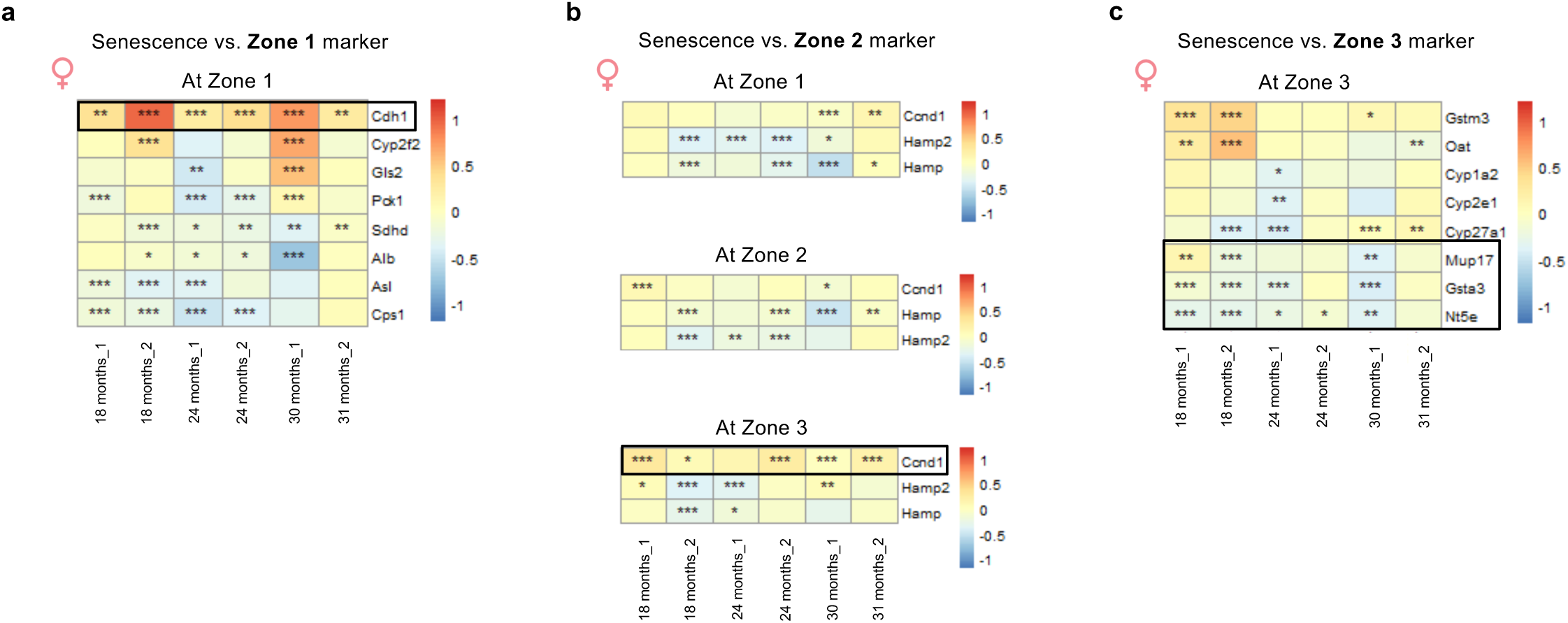
Correlation of the high-senescence-score-pixels with the expression of zonation markers in WT female mouse livers (related to Figure 4) a. Heatmap showing the correlation between high-senescence-score-pixels and the expression of zone 1 markers (y-axis) within zone 1 of each WT female mouse liver sample at different ages (x-axis). Correlation coefficient ranging from −1.0 (perfect negative relationship, blue) to +1.0 (perfect positive relationship, red). T-distribution test: \**p* < 0.05, \*\**p* < 0.01, \*\*\**p* < 0.001. b. Heatmap showing the correlation between high-senescence-score-pixels and the expression of zone 2 markers (y-axis) across three liver zones of each WT female mouse liver sample at different ages (x-axis). Correlation coefficient ranging from −1.0 (perfect negative relationship, blue) to +1.0 (perfect positive relationship, red). T-distribution test: \**p* < 0.05, \*\**p* < 0.01, \*\*\**p* < 0.001. Across all three hepatic zones, zone 2 marker *Ccnd1* shows the strongest correlation with senescence at zone 3 regions. c. Heatmap showing the correlation between high-senescence-score-pixels and the expression of zone 3 markers (y-axis) within zone 3 of each WT female mouse liver sample at different ages (x-axis). Correlation coefficient ranging from −1.0 (perfect negative relationship, blue) to +1.0 (perfect positive relationship, red). T-distribution test: \**p* < 0.05, \*\**p* < 0.01, \*\*\**p* < 0.001.

**Extended Data Figure 5.**
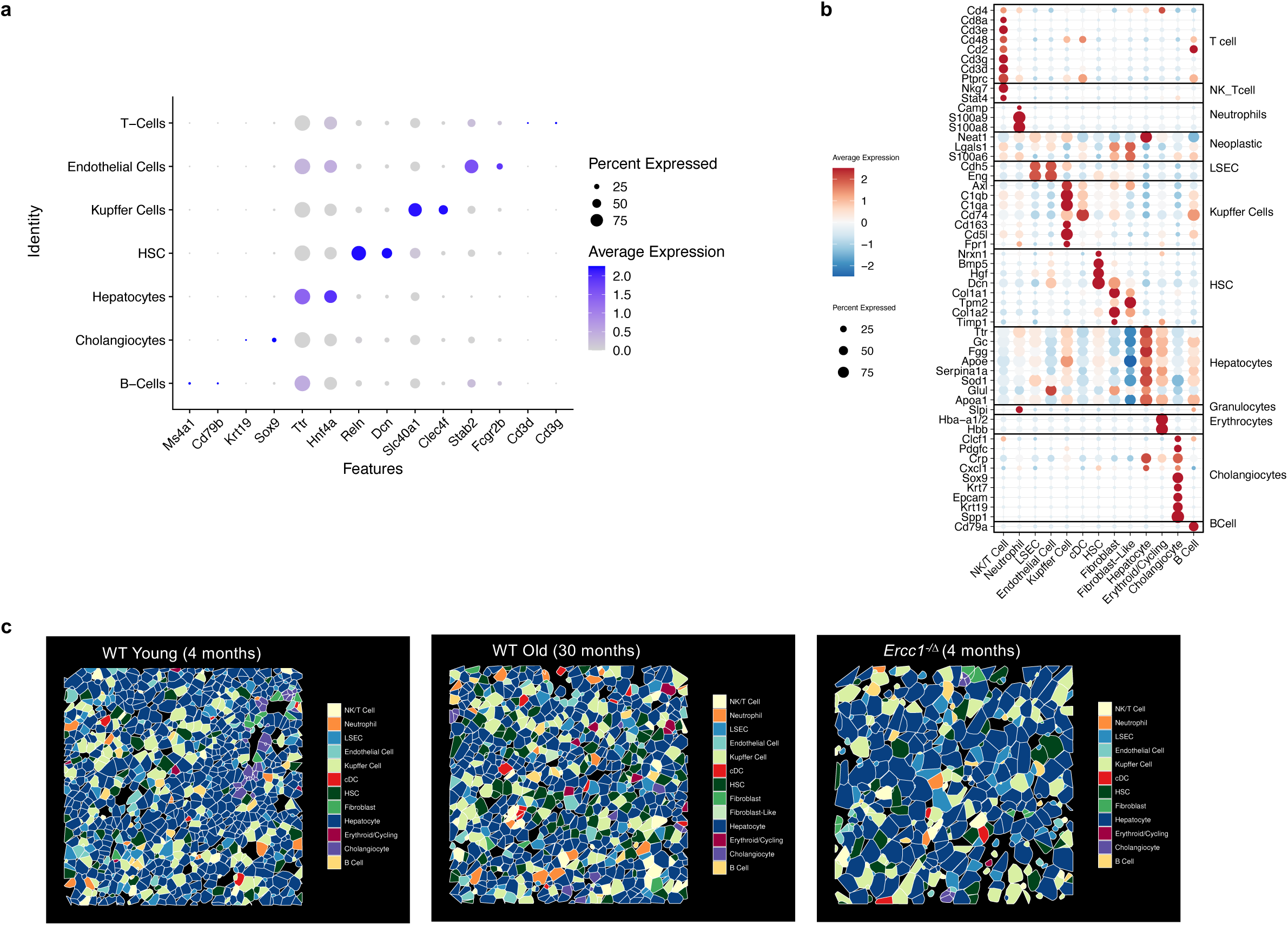
Identification of senescent cells in aged mouse livers by snRNA-seq and CosMx spatial transcriptomics (related to Figures 6) a. Gene panel (x-axis) used to identify cell types (y-axis) in murine livers from snRNA-seq data. b. Gene panel (y-axis) used to identify cell types (x-axis) in murine livers from CosMx transcriptomics data. c. Representative images of cell segmentation of CosMx data by cell types in young (4-month), old (30-month) WT and *Ercc1^−/Δ^* (4-month). Each image represents one full field of view (FOV) of 511 x 511 µM^2^.

**Extended Data Figure 6.**
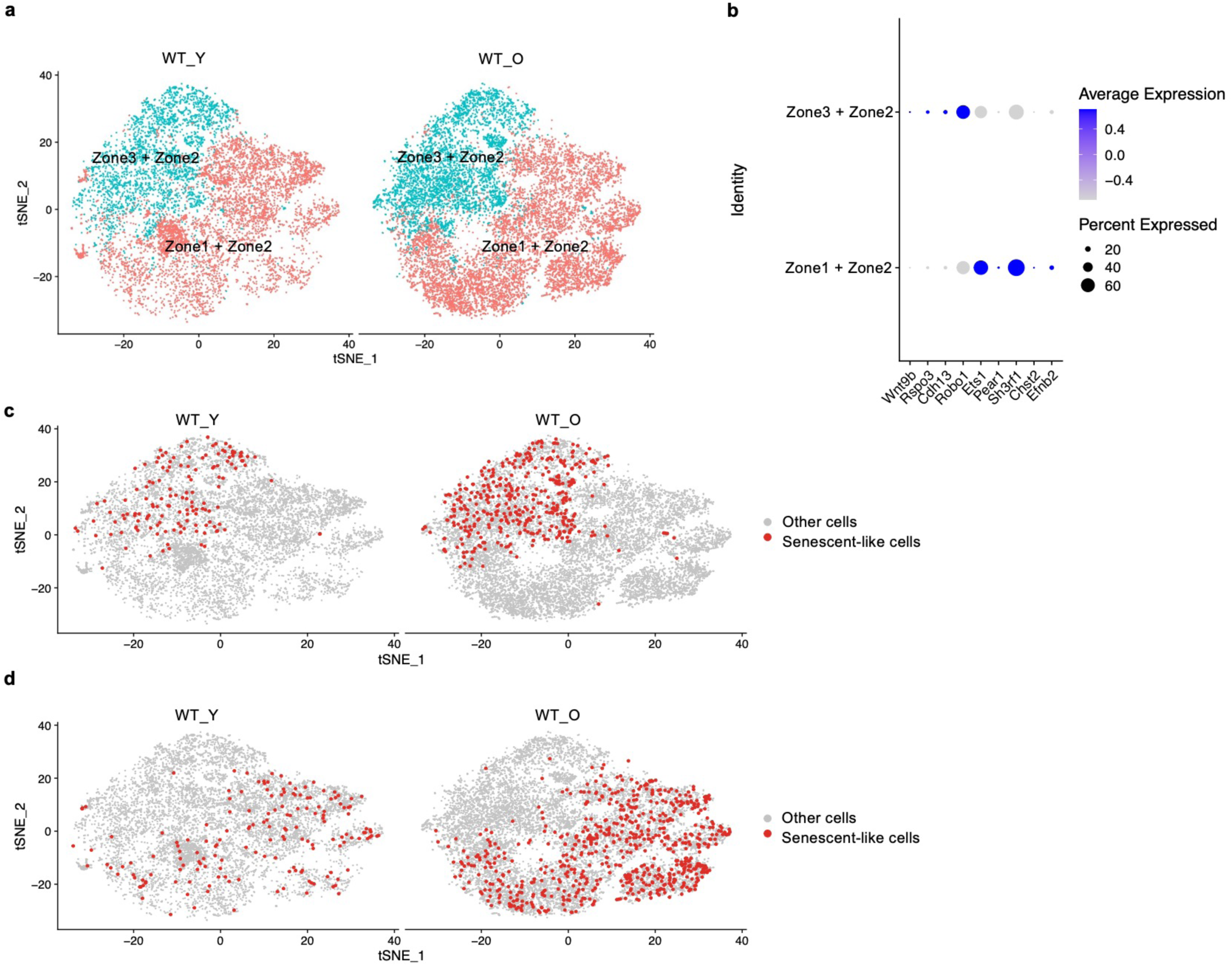
snRNA-seq studies revealed senescence-associated signatures in aged endothelial cells (related to Figure 6) a. tSNE plot showing the identification of endothelial cell clusters by hepatic zone in snRNA-seq data from young and old WT mice, described in Figure 6b. b. Gene panel (x-axis) used to identify endothelial cells by liver zone (y-axis) in snRNA-seq data. c. tSNE plot showing senescent-like endothelial cells in pericentral + midlobular zones (zone 3 + zone 2) based on differential expression of senescence-associated genes (SASP, cell cycle inhibitor, and anti-apoptosis) between young and old WT mice. d. tSNE plot showing senescent-like endothelial cells in periportal + midlobular zones (zone 1 + zone 2) based on differential expression of senescence-associated genes (SASP, cell cycle inhibitor, and anti-apoptosis) between young and old WT mice.

**Extended Data Figure 7.**
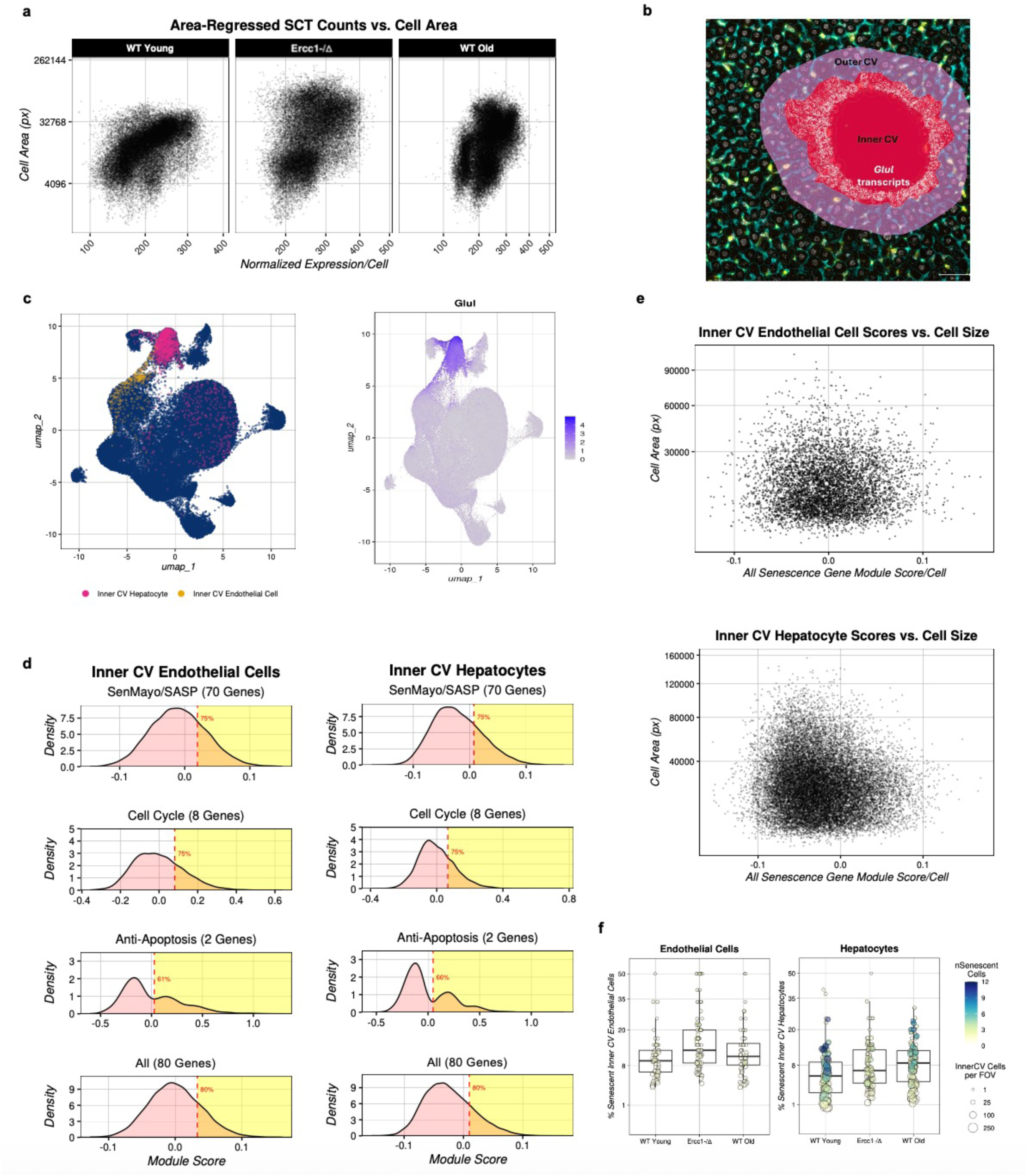
Senescent cell identification in old WT and *Ercc1*^−/Δ^ mouse livers by CosMx spatial transcriptomics (related to Figures 6 and 7) a. Scatter plots showing area regression of total transcript counts after controlling for read depth variation based on cell size for CosMx datasets. WT young (∼ 4 months) n=3, WT old (∼ 30 months) n=3, *Ercc1*^−/Δ^ young (4 months) n=4, all female mice. Px stands for pixels. SCT stands for SCTransform-normalized counts. b. Representative image showing manual annotation of the inner and outer central vein (CV) area in mouse liver, using expression of the pericentral zone marker gene *Glul* (white) as guide, with inner CV identified as the cells with strong *Glul* signals (usually the 1-2 layers surrounding the central vein), and outer CV identified as roughly another 3 layers of hepatocytes adjacent to *Glul* positive cells. c. UMAP plot showing one distinct subcluster of inner CV endothelial cells (in gold color) and one distinct subcluster of inner CV hepatocytes (in magenta color), the latter of which overlapped with high *Glul* gene expression shown in the right panel. d. Distribution plots showing the scoring strategy used to identify senescent endothelial cells (left) and hepatocytes (right) in the inner CV regions. The cutoff for calling a cell senescent is indicated with a red vertical bar in each panel. e. Scatter plots showing area regression of total transcripts counts of the 80 senescence-associated genes after controlling for read depth variation based on cell size for endothelial cells (top) and hepatocytes (bottom) in the inner CV regions. Px stands for pixels. f. Box-whisker plots showing the percentage of endothelial cells and hepatocytes in the inner CV regions that are senescent based on the strategy illustrated in d. The mean percentage of senescent endothelial cells (left) or hepatocytes (right) is indicated by the horizontal bars. Each circle represents a field of view (FOV) and the size of the circle indicates the number of cells in that FOV.

**Extended Data Figure 8.**
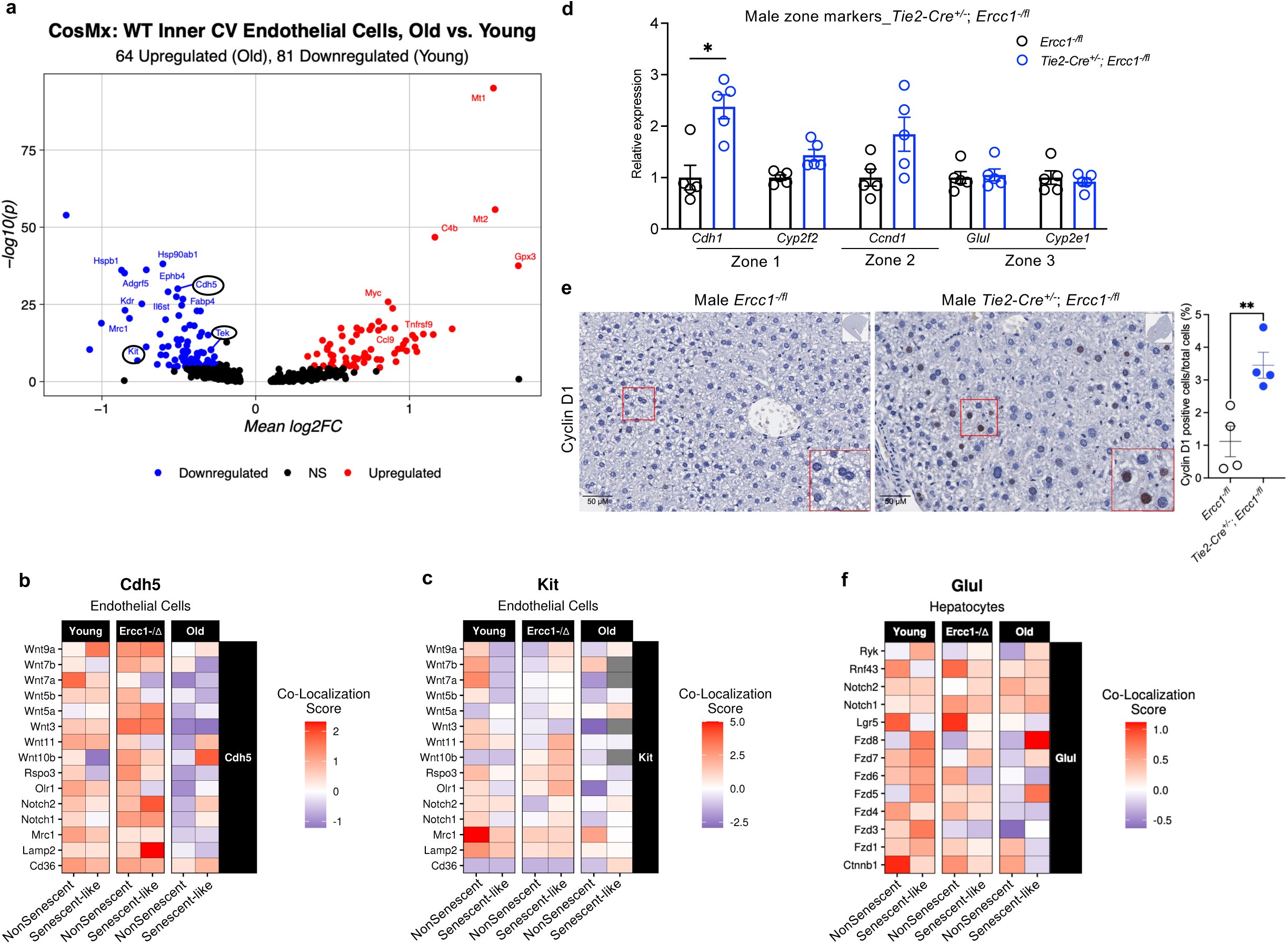
Loss of cell identity in senescent endothelial cells and hepatocytes with disrupted WNT signaling contributed to zonation changes (related to Figure 6 and 7) a. Volcano plot of differentially expressed genes (DEGs) in endothelial cells in the inner central vein (CV) region from old (30-month-old) vs. young (4-month-old) WT mouse livers identified by CosMx spatial transcriptomics. WT young n=3 mice, WT old n=3 mice. Notably, cell identity markers are down-regulated (black circles). b. Co-localization of Wnt signaling pathway gene expression in senescent vs. non-senescent *Cdh5^+^* inner CV endothelial cells of young WT, *Ercc1^−/Δ^* (4-month-old), and old WT mouse livers identified by CosMx spatial transcriptomics. WT young n=3 mice, *Ercc1*^−/Δ^ n=4 mice, WT old n=3 mice. c. Same as b except for inner CV endothelial cells identified by *Kit^+^* expression. d. qPCR measurement of expression of zone marker gene expression in *Tie2-Cre^+/−^*;*Ercc1^−/fl^*and control male mouse livers. Each circle represents an individual mouse (n=5 per group). Multiple unpaired *t*-tests and adjusted *p* value is presented as: \**p* < 0.05. e. Representative images of immunohistochemical staining of midlobular zone marker Cyclin D1 in control and *Tie2-Cre^+/−^*;*Ercc1^−/fl^*male mouse livers. Two-tailed student *t*-test was applied for Cyclin D1 positive cell (%) quantification comparison, \*\**p* < 0.01. n=4 for *Ercc1^−/fl^*(18 - 20 weeks), n=4 for *Tie2-Cre^+/−^*;*Ercc1^−/fl^*(16 - 20 weeks) male mice. f. Same as b and c except for inner CV hepatocytes identified by *Glul^+^* expression.

**Extended Data Figure 9.**
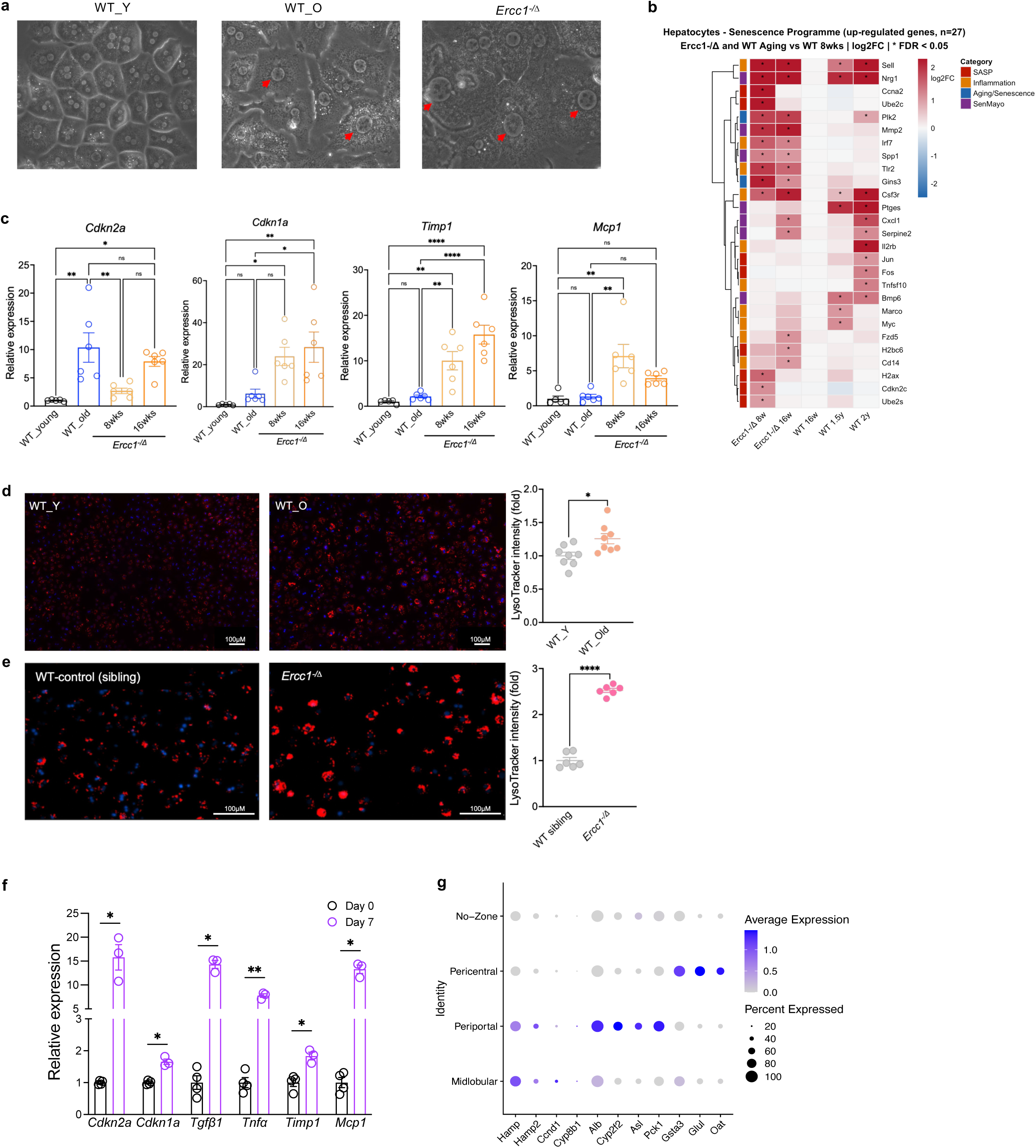
Senescent hepatocytes characterization from old WT and *Ercc1*^−/Δ^ mouse livers (related to Figures 7 and 8) a. Representative images of hepatocytes isolated from young WT, old WT and *Ercc1*^−/Δ^ mice after 24 hours in culture (20x magnification) illustrating cell hypertrophy and nuclear polyploidy (red arrows) in cells from old or progeroid mice. b. Heatmap of the significantly up-regulated expression of SASP, inflammation, aging/senescence and SenMayo genes (y-axis) identified in WT and *Ercc1*^−Δ^ primary hepatocytes relative to 8-week-old WT mice in bulk RNA-seq data, *FDR < 0.05. c. qPCR measurement of the relative expression of senescence-associated genes in primary hepatocytes isolated from old WT (at age >2-years-old) and *Ercc1*^−/Δ^ mice (at age of 8-weeks and 16-weeks-old) relative to young WT (at age of 8-weeks-old). Each circle represents one sample isolated from an individual mouse liver (n=5-6 per group). One-way ANOVA analysis and adjusted *p* values are presented as: \**p* < 0.05, \*\**p* < 0.01, \*\*\*\**p* < 0.0001. d. Lysotracker staining of primary hepatocytes isolated from young and old WT mice, quantitated on the right, where each dot represents a technical replicate of the hepatocytes isolated from one animal. Two-tailed student *t*-test and *p* value is presented: \**p* < 0.05. Experiments were repeated at least twice. e. Same as d. except for *Ercc1*^−/Δ^ mice, \*\*\*\**p* < 0.0001. Experiments were repeated at least three times. f. qPCR measurement of the expression of senescence-associated genes in primary hepatocytes after 7 days in culture at 20% O_2_. Each dot represents a technical replicate of the hepatocytes isolated from a 12-week-old mouse liver, relative to cells freshly isolated (Day 0) from the same mouse liver. Multiple unpaired *t*-tests and adjust *p* values are presented as: \**p* < 0.05, \*\**p* < 0.01. Experiments were repeated at least 3 times. g. Gene panel (x-axis) used to identify the different zone of the hepatocytes analyzed by snRNA-seq data.

## References

1. Anantharaju, A., A. Feller, and A. Chedid, Aging Liver. A review. Gerontology, 2002. 48(6): p. 343–53.

2. Yao, Y.Q., Q.Y. Cao, and Z. Li, Delaying liver aging: Analysis of structural and functional alterations. World J Gastroenterol, 2025. 31(15): p. 103773.

3. Nikopoulou, C., et al., Spatial and single-cell profiling of the metabolome, transcriptome and epigenome of the aging mouse liver. Nat Aging, 2023. 3(11): p. 1430–1445.

4. Zhao, H., et al., Identifying specific functional roles for senescence across cell types. Cell, 2024. 187(25): p. 7314–7334 e21.

5. Idda, M.L., et al., Survey of senescent cell markers with age in human tissues. Aging (Albany NY), 2020. 12(5): p. 4052–4066.

6. Bird, T.G., et al., TGFbeta inhibition restores a regenerative response in acute liver injury by suppressing paracrine senescence. Sci Transl Med, 2018. 10(454).

7. Sanfeliu-Redondo, D., A. Gibert-Ramos, and J. Gracia-Sancho, Cell senescence in liver diseases: pathological mechanism and theranostic opportunity. Nat Rev Gastroenterol Hepatol, 2024. 21(7): p. 477–492.

8. Khosla, S., et al., The role of cellular senescence in ageing and endocrine disease. Nat Rev Endocrinol, 2020. 16(5): p. 263–275.

9. Ogrodnik, M., et al., Cellular senescence drives age-dependent hepatic steatosis. Nat Commun, 2017. 8: p. 15691.

10. Grosse, L., et al., Defined p16(High) Senescent Cell Types Are Indispensable for Mouse Healthspan. Cell Metab, 2020. 32(1): p. 87–99 e6.

11. Krizhanovsky, V., et al., Senescence of activated stellate cells limits liver fibrosis. Cell, 2008. 134(4): p. 657–67.

12. Moncsek, A., et al., Targeting senescent cholangiocytes and activated fibroblasts with B-cell lymphoma-extra large inhibitors ameliorates fibrosis in multidrug resistance 2 gene knockout (Mdr2(−/−)) mice. Hepatology, 2018. 67(1): p. 247–259.

13. Cazzagon, N., et al., Cholangiocyte senescence in primary sclerosing cholangitis is associated with disease severity and prognosis. JHEP Rep, 2021. 3(3): p. 100286.

14. Salladay-Perez, I.A., et al., p21(+)TREM2(+) senescent macrophages fuel inflammaging and metabolic dysfunction-associated steatotic liver disease. Nat Aging, 2026.

15. Cunningham, R.P. and N. Porat-Shliom, Liver Zonation - Revisiting Old Questions With New Technologies. Front Physiol, 2021. 12: p. 732929.

16. Raven, A., et al., Hepatic zonation determines tumorigenic potential of mutant beta-catenin. Nature, 2025.

17. Paolini, E., et al., Hepatic Zonation in MASLD: Old Question, New Challenge in the Era of Spatial Omics. Int J Mol Sci, 2025. 26(21).

18. Torre, C., C. Perret, and S. Colnot, Transcription dynamics in a physiological process: beta-catenin signaling directs liver metabolic zonation. Int J Biochem Cell Biol, 2011. 43(2): p. 271–8.

19. Burke, Z.D., et al., Liver zonation occurs through a beta-catenin-dependent, c-Myc-independent mechanism. Gastroenterology, 2009. 136(7): p. 2316–2324 e1-3.

20. Benhamouche, S., et al., Apc tumor suppressor gene is the “zonation-keeper” of mouse liver. Dev Cell, 2006. 10(6): p. 759–70.

21. Planas-Paz, L., et al., The RSPO-LGR4/5-ZNRF3/RNF43 module controls liver zonation and size. Nat Cell Biol, 2016. 18(5): p. 467–79.

22. Annunziato, S., T. Sun, and J.S. Tchorz, The RSPO-LGR4/5-ZNRF3/RNF43 module in liver homeostasis, regeneration, and disease. Hepatology, 2022. 76(3): p. 888–899.

23. Duan, J.L., et al., Notch-Regulated c-Kit-Positive Liver Sinusoidal Endothelial Cells Contribute to Liver Zonation and Regeneration. Cell Mol Gastroenterol Hepatol, 2022. 13(6): p. 1741–1756.

24. He, B., et al., Spatial regulation of glucose and lipid metabolism by hepatic insulin signaling. Cell Metab, 2025. 37(7): p. 1568–1583 e7.

25. Hu, S., et al., Single-cell spatial transcriptomics reveals a dynamic control of metabolic zonation and liver regeneration by endothelial cell Wnt2 and Wnt9b. Cell Rep Med, 2022. 3(10): p. 100754.

26. He, L., et al., Proliferation tracing reveals regional hepatocyte generation in liver homeostasis and repair. Science, 2021. 371(6532).

27. Wei, Y., et al., Liver homeostasis is maintained by midlobular zone 2 hepatocytes. Science, 2021. 371(6532).

28. Zhang, Y., et al., Zonated mechanosensing by PIEZO1 controls liver regeneration. Science, 2026. 393(6806): p. eaef0825.

29. Stavropoulos, A., et al., Coordinated activation of TGF-beta and BMP pathways promotes autophagy and limits liver injury after acetaminophen intoxication. Sci Signal, 2022. 15(740): p. eabn4395.

30. Herrera, B., A. Addante, and A. Sanchez, BMP Signalling at the Crossroad of Liver Fibrosis and Regeneration. Int J Mol Sci, 2017. 19(1).

31. Viatte, L. and S. Vaulont, Hepcidin, the iron watcher. Biochimie, 2009. 91(10): p. 1223–8.

32. Saul, D., et al., A new gene set identifies senescent cells and predicts senescence-associated pathways across tissues. Nat Commun, 2022. 13(1): p. 4827.

33. Yousefzadeh, M.J., et al., Tissue specificity of senescent cell accumulation during physiologic and accelerated aging of mice. Aging Cell, 2020. 19(3): p. e13094.

34. Sinha, S., et al., Aging disrupts hepatocyte zonation homeostasis in mice and humans. Hepatology, 2025.

35. Yang, Q., et al.,, Single-cell analysis reveals altered liver zonation in aging-associated insulin resistance. Metabolism Clinical and Experimental, 2026. 177.

36. Chong, M., et al., CD36 initiates the secretory phenotype during the establishment of cellular senescence. EMBO Rep, 2018. 19(6).

37. Carpintero-Fernandez, P., et al., Genome wide CRISPR/Cas9 screen identifies the coagulation factor IX (F9) as a regulator of senescence. Cell Death Dis, 2022. 13(2): p. 163.

38. Gregg, S.Q., et al., A mouse model of accelerated liver aging caused by a defect in DNA repair. Hepatology, 2012. 55(2): p. 609–21.

39. Valle-Encinas, E. and T.C. Dale, Wnt ligand and receptor patterning in the liver. Curr Opin Cell Biol, 2020. 62: p. 17–25.

40. Canal, F., et al., Generation of Mice with Hepatocyte-Specific Conditional Deletion of Notum. PLoS One, 2016. 11(3): p. e0150997.

41. Ovejero, C., et al., Identification of the leukocyte cell-derived chemotaxin 2 as a direct target gene of beta-catenin in the liver. Hepatology, 2004. 40(1): p. 167–76.

42. Kisanuki, Y.Y., et al., Tie2-Cre transgenic mice: a new model for endothelial cell-lineage analysis in vivo. Dev Biol, 2001. 230(2): p. 230–42.

43. Wang, Z. and P.A. Burke, Hepatocyte nuclear factor-4alpha interacts with other hepatocyte nuclear factors in regulating transthyretin gene expression. FEBS J, 2010. 277(19): p. 4066–75.

44. Rufibach, L.E., et al., Transcriptional regulation of the human hepatic lipase (LIPC) gene promoter. J Lipid Res, 2006. 47(7): p. 1463–77.

45. Walesky, C., et al., Hepatocyte-specific deletion of hepatocyte nuclear factor-4alpha in adult mice results in increased hepatocyte proliferation. Am J Physiol Gastrointest Liver Physiol, 2013. 304(1): p. G26–37.

46. Lin, Y., et al., Bmal1 regulates circadian expression of cytochrome P450 3a11 and drug metabolism in mice. Commun Biol, 2019. 2: p. 378.

47. Tan, X., et al., Conditional deletion of beta-catenin reveals its role in liver growth and regeneration. Gastroenterology, 2006. 131(5): p. 1561–72.

48. Tan, X., et al., Beta-catenin deletion in hepatoblasts disrupts hepatic morphogenesis and survival during mouse development. Hepatology, 2008. 47(5): p. 1667–79.

49. Decaens, T., et al., Stabilization of beta-catenin affects mouse embryonic liver growth and hepatoblast fate. Hepatology, 2008. 47(1): p. 247–58.

50. Nemeth, E., et al., Hepcidin regulates cellular iron efflux by binding to ferroportin and inducing its internalization. Science, 2004. 306(5704): p. 2090-3.

51. Canali, S., et al., Endothelial cells produce bone morphogenetic protein 6 required for iron homeostasis in mice. Blood, 2017. 129(4): p. 405–414.

52. Chen, S., et al., Transforming Growth Factor beta1 (TGF-beta1) Activates Hepcidin mRNA Expression in Hepatocytes. J Biol Chem, 2016. 291(25): p. 13160–74.

53. Siraj, Y., et al., IGFBP7 is a key component of the senescence-associated secretory phenotype (SASP) that induces senescence in healthy cells by modulating the insulin, IGF, and activin A pathways. Cell Commun Signal, 2024. 22(1): p. 540.

54. Wang, Y., et al., Depletion of Igfbp7 alleviates zebrafish NAFLD progression through inhibiting hepatic ferroptosis. Life Sci, 2023. 332: p. 122086.

55. Andrews, T.S., et al., Single-Cell, Single-Nucleus, and Spatial RNA Sequencing of the Human Liver Identifies Cholangiocyte and Mesenchymal Heterogeneity. Hepatol Commun, 2022. 6(4): p. 821–840.

56. Niedernhofer, L.J., et al., A new progeroid syndrome reveals that genotoxic stress suppresses the somatotroph axis. Nature, 2006. 444(7122): p. 1038–43.

57. Xu, J., et al., Impaired iron status in aging research. Int J Mol Sci, 2012. 13(2): p. 2368–2386.

58. Hahn, P., et al., Age-dependent and gender-specific changes in mouse tissue iron by strain. Exp Gerontol, 2009. 44(9): p. 594–600.

59. Liu, J., et al., Iron accumulation with age alters metabolic pattern and circadian clock gene expression through the reduction of AMP-modulated histone methylation. J Biol Chem, 2022. 298(6): p. 101968.

60. Park, S.H., et al., Effects of Aging on the Severity of Liver Injury in Mice With Iron Overload. J Gastroenterol Hepatol, 2025. 40(4): p. 1016–1025.

61. Milic, S., et al., The Role of Iron and Iron Overload in Chronic Liver Disease. Med Sci Monit, 2016. 22: p. 2144–51.

62. Zhang, D., et al., Inferring super-resolution tissue architecture by integrating spatial transcriptomics with histology. Nat Biotechnol, 2024. 42(9): p. 1372–1377.

63. Ritchie, M.E., et al., limma powers differential expression analyses for RNA-sequencing and microarray studies. Nucleic Acids Res, 2015. 43(7): p. e47.

64. Hao, Y., et al., Integrated analysis of multimodal single-cell data. Cell, 2021. 184(13): p. 3573–3587 e29.

65. Korsunsky, I., et al., Fast, sensitive and accurate integration of single-cell data with Harmony. Nat Methods, 2019. 16(12): p. 1289–1296.

